# Longitudinal genomic surveillance of a UK intensive care unit shows a lack of patient colonisation by multi-drug resistant Gram-negative pathogens

**DOI:** 10.1101/2024.07.25.605077

**Authors:** Ann E. Snaith, Robert A. Moran, Rebecca J. Hall, Anna Casey, Liz Ratcliffe, Willem van Schaik, Tony Whitehouse, Alan McNally

## Abstract

Vulnerable patients in an intensive care unit (ICU) setting are at high risk of infection from bacteria including gut-colonising *Escherichia coli* and *Klebsiella* species. Complex ICU procedures often depend on successful antimicrobial treatment, underscoring the importance of understanding the extent of patient colonisation by multi-drug resistant organisms (MDRO) in large UK ICUs. Previous work on ICUs globally uncovered high rates of colonisation by and transmission of MDRO, but the situation in UK ICUs is less understood.

Here, we investigated the diversity and antibiotic resistance gene (ARG) carriage of bacteria present in one of the largest UK ICUs at the Queen Elizabeth Hospital Birmingham (QEHB), focussing primarily on *E. coli* as both a widespread commensal and a globally disseminated multidrug resistant pathogen. Samples were taken during highly restrictive COVID-19 control measures from May – December 2021. Whole-genome and metagenomic sequencing were used to detect and report strain level colonisation of patients, focussing on *E. coli* sequence types (STs), their colonisation dynamics, and antimicrobial resistance (AMR) gene carriage.

We found a lack of multidrug resistance (MDR) in the QEHB. Only one carbapenemase-producing organism was isolated, a *Citrobacter* carrying *bla*_KPC-2_. There was no evidence supporting the spread of this strain, and there was little evidence overall of nosocomial acquisition or circulation of colonising *E. coli*. Whilst 22 different *E. coli* STs were identified, only one strain of the pandemic ST131 lineage was isolated. This ST131 strain was non-MDR and was found to be a clade A strain, associated with low levels of antibiotic resistance. Overall, the QEHB ICU had very low levels of pandemic or MDR strains, a result which may be influenced in part by the strict COVID-19 control measures in place at the time. Employing some of these infection prevention and control measures where reasonable in all ICUs might therefore assist in maintaining low levels of nosocomial MDR.

**Impact statement:** This study contributes to current literature on the potential routes for AMR spread in a healthcare setting. This study used whole genome sequencing (WGS) to investigate at strain-level bacterial species (including *E. coli)* colonising the gut of long-stay patients in the ICU. WGS in combination with patient ward movement and prescribing information was used to assess any links or driving factors in strain acquisition and AMR spread in the ICU.

Our study gives an insight at a point in time where infection and prevention control restrictions and awareness were high due to the COVID-19 pandemic, combined with local and national travel restrictions and isolation criteria. Consequently, it provides a novel longitudinal dataset that gives a picture of colonising *E. coli* in a sheltered ICU patient population. Trends seen in this *E. coli* population are likely linked to the United Kingdom in 2021 rather than the global picture that may have been seen prior to the COVID-19 pandemic.

**Data summary:** All supporting data, code and protocols have been provided within the article or through supplementary data files. All genomic and metagenomic data are available from NCBI under BioProject accession PRJNA1136496.

All relevant metadata is provided in supplementary data files.

## Introduction

The increasing incidence of antimicrobial resistance (AMR) internationally has a significant impact in the healthcare setting, causing increased morbidity and mortality in both the immunocompetent and immunocompromised (1). Major surgical procedures and other interventions depend on antimicrobials to protect high risk intensive care unit (ICU) patients from infection and assist their recovery (2). Gut colonisation is one potential source of infection in vulnerable patients in an ICU and consequently potential AMR transmission. Multidrug resistant organisms (MDROs) frequently carry a variety of virulence factors which help them to easily colonise the gut (3, 4) and with the potential to be a risk of infection in this vulnerable population.

Recent studies in South East Asia and China have highlighted significant transmission of MDROs (including carbapenem resistant *Escherichia coli, Klebsiella pneumoniae* and *Acinetobacter baumannii*) at high levels in the ICU environment (5-9). In some cases this transmission has been attributed to breaches of infection prevention and control (IPC) measures. Travel or inpatient hospital stays in endemic areas (e.g., S. E. Asia) increases chances of gut colonisation with these MDROs (10, 11). Colonisation with MDROs is a particular risk for ICU patients who generally have longer hospital stays and may be subject to a wide range of interventions (e.g., mechanical ventilators, intravenous lines), which can provide other opportunities for colonisation and subsequent infection (12). These patients frequently have a weakened immune response, chronic pathologies or trauma, so often receive many medications (e.g., painkillers, steroids) and it is more likely that antimicrobials will be prescribed in this environment (13, 14). Increased use of antimicrobials can select for drug resistant isolates present in the hospital environment and drive the development of MDR. Long stays in hospital environments can lead to colonisation with organisms not frequently seen in healthy patients (e.g., *Acinetobacter*) and disturbed microbiomes with overgrowth of commensal species (e.g., *Candida, E. coli*) (15, 16). Bacterial isolates found colonising ICUs are known to differ globally especially in the different burdens and profiles of AMR found (17-21). As recent studies have shown, *E. coli* and *Klebsiella* species are commonly seen in ICUs, both in the UK and globally (18, 22).

A prospective observational shotgun sequencing study on 24 QEHB patients in 2017-2018 investigated changes in the patient gut microbiome during their ICU stay (15). As anticipated during an ICU stay, gut microbiome diversity decreased and the commensal gut flora changed, with overgrowth of *Candida* species, *Enterococcus faecium* and *E. coli* (15). Similarly, other ICU studies have found dysbiosis of patient gut microbiome and reduced microbiota diversity after hospital stays with an increased likelihood of serious nosocomial infections (e.g., sepsis) (23, 24). In contrast to patients on a general ward, ICU patients have been found to have a significantly increased risk of colonisation by an antimicrobial resistant organism (e.g., ampicillin and/or cephalosporin-resistant *E. coli*), demonstrating the effect of the difference in environment, severity of illness and extent of treatment experienced by ICU patients (20). With this study (IMPACT2) we aimed to explore the diversity and levels of gut colonising bacteria in high-risk patients in the UK’s largest ICU, focussing primarily on *E. coli* and the prevalence of MDR. This study coincided with strict IPC protocols enforced during the COVID-19 pandemic, providing an ideal opportunity to monitor how these procedures might affect the transmission of MDR.

## Methods

### Sample Collection

All bacterial isolates in the IMPACT2 study were obtained from the intensive care unit (ICU) at Queen Elizabeth Hospital Birmingham (QEHB), University Hospitals Birmingham, NHS Foundation Trust (UHB). QEHB houses the largest ICU in the UK by bed-base (68 funded beds) and annual patient throughput (approximately 5000 patients per annum), which is divided into four ICU wards by specialty (liver/specialty ICU, general/trauma/burns ICU, neurology and neurosurgical ICU, and cardiac ICU) and also has 15 side rooms. The trial was approved by the Yorkshire & The Humber - Leeds East Research Ethics Committee (Reference, 20/YH/0067). Study inclusion criteria were that patients must be aged over 18 years and admitted to the ICU with the expectation they would remain there for greater than 48 hours. Study exclusion criteria included patients aged under 18 years, patients admitted to the ICU who were not expected to remain there for greater than 48 hours and patients who had opened their bowels whilst on ICU before enrolment to this study. Patients were recruited and consented to the study by QEHB research nurses after discussion with an ICU consultant about the likelihood they would remain on the ward for over 48 hours. The first stool sample passed by the patient was stored. Patients then remained in the study for the duration of their stay and their stool samples were collected by trained research nurses until discharge or patient death on the ICU. Metadata was collected including sex, age, admission date to ward, departure date from ward, bed history, microbiology diagnostic test results and antibiotic prescribing history.

### Stool sample Processing

Bacterial isolates were selected for whole genome sequencing (WGS) using bacterial culture methods on stool culture. In order to isolate colonies of interest 100 μg of each stool sample was incubated in 10 mL of LB (Miller) broth overnight (shaking 200 rpm, 37 °C). Overnight stool broths were then diluted in phosphate buffered saline (PBS) to concentrations ranging from 10^−1^ to 10^−6^. Initially 100 μL from each dilution (from neat to 10^−6^) was plated on both MacConkey agar (Sigma) and ESBL Chromoselect agar (Sigma). This range of dilutions was reduced to neat, 10^−1^, 10^−3^ and 10^−6^ over the course of the study. Dilution plates of MacConkey and ESBL agar were both incubated overnight at 37 °C and representative populations of all bacterial isolates were stored in 25% glycerol. Single isolates from each sample of *E. coli* and *Klebsiella* were selected for short read WGS by MicrobesNG.

### Bioinformatics

All short read sequencing data (MicrobesNG) were provided as read files that had undergone initial trimming (Trimmomatic [v0.3]) with a sliding window quality of Q15 to remove adapters. Genome assemblies for all isolates were created from their trimmed read files using the de Novo assembler SPAdes (v3.13.0) (25). All assemblies were annotated using Prokka (v1.14.6) (26). *E. coli* isolates were allocated to a phylogroup using ClermonTyping (27). Sequence data was analysed using mlst (https://github.com/tseemann/mlst) (v2.23) to determine species and sequence types. *Klebsiella* isolates were run through Kleborate (v2.2.0) for speciation and clone typing. Kleborate output files were run through the Kleborate-viz web app (https://kleborate.erc.monash.edu/) (28). Assemblies were analysed using ABRicate (v1.0.1) with the ResFinder (29) and PlasmidFinder (30) databases to detect acquired AMR genes and plasmid replicons. Virulence genes were identified using ABRicate (v0.4.8) and the vfdb virulence factor database (31). Core gene alignments were created using Panaroo (v1.2.10). Snippy (v4.6.0) was used to call single nucleotide polymorphism (SNP)s between the reference Prokka annotated GenBank (gbk) file and isolates of the same ST. High-resolution single nucleotide polymorphism analysis used the first isolate of strain as a reference to conduct strain sharing dynamic analysis. Core SNP alignments were generated using from Snippy-core (v4.6.0). SNP distances for all species were calculated using snp-dists (v0.8.2). Within participant or patient comparisons used the earliest isolate as the reference. All further phylogenetic trees for all datasets were constructed using IQTREE (v2.0.3) on either Panaroo or Snippy core SNP alignment files.

Metagenomic reads were screened for antibiotic resistance genes using ShortBRED (32) with the CARD database(33).

## Results

### An overview of the study patient population

Fifty-seven patients were recruited to this study (IMPACT2), with our investigation focussed on the 44 ICU patients recruited between May 2021 and December 2021. Twenty-one patients were excluded as they either did not produce stool samples during their ICU admission or withdrew from the study, leaving a final total of twenty-three patients (Table S1). Overall, 188 stool samples from 23 patients were processed between May 2021 and December 2021 (Figure 1).

**Fig. 1.**
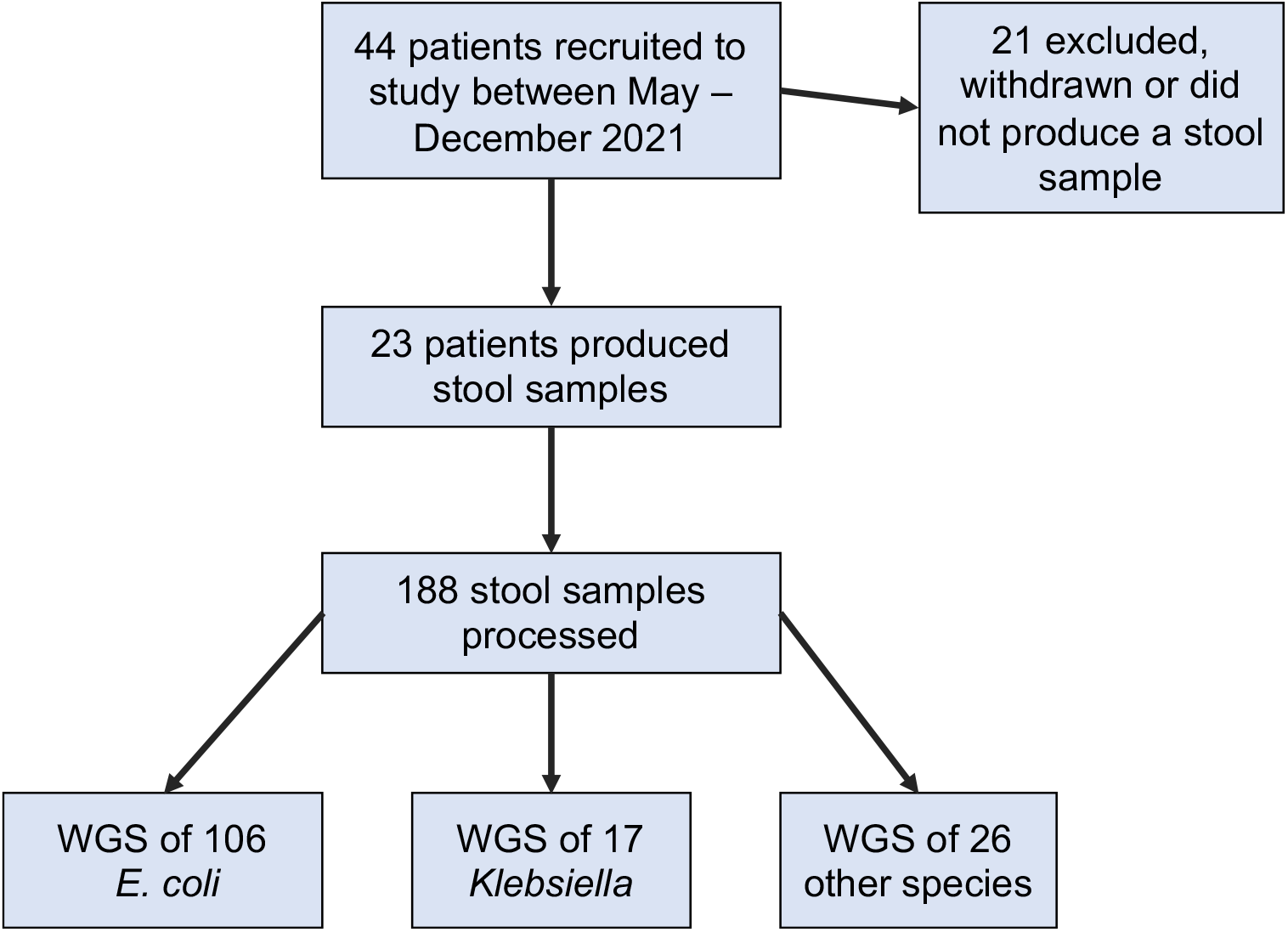
Workflow of the study showing the number of patients recruited and the number of isolates whole genome sequenced.

Inpatient stays ranged from three to 55 days. In total, 149 putative *E. coli* and *Klebsiella* isolates were whole genome sequenced (Figure 1). Of the 23 patients that produced stool samples, 18 were colonised with *E. coli* at some point during their ICU stay and six were colonised with *Klebsiella*. Patients, ranging in age between 20 and 80 years old, stayed on the ICU for a mean of seven days (three – 14 days) before they produced a stool sample. Reasons for ICU admission varied, including stays following planned elective surgery or as a result of other trauma or need for critical care (Table 1).

**Table 1.**
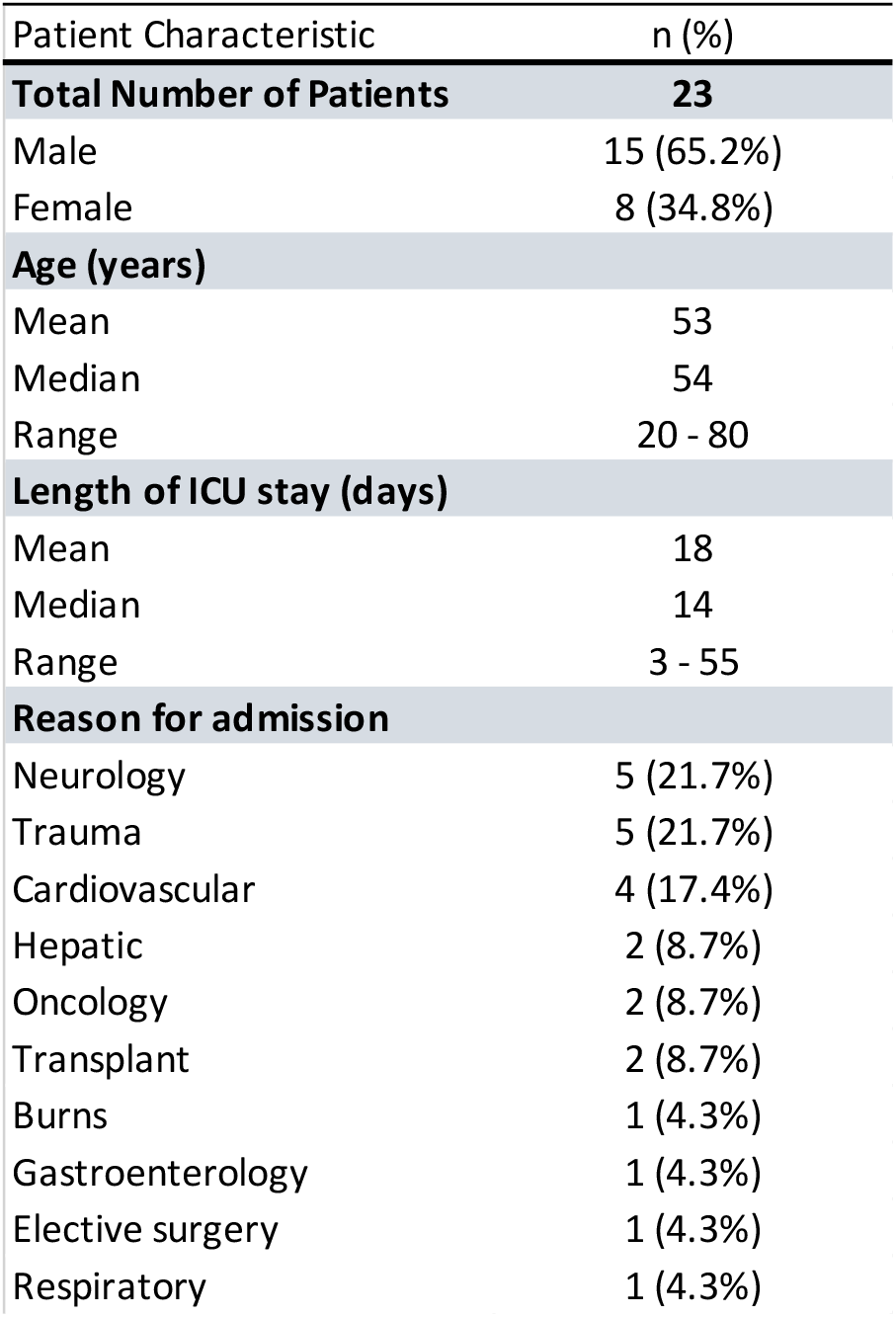
Characteristics of patients included in the analysis. Further detail provided in Supplementary File S1.

| Patient Characteristic | n (%) |
| --- | --- |
| **Total Number of Patients** | **23** |
| Male | 15 (65.2%) |
| Female | 8 (34.8%) |
| **Age (years)** |  |
| Mean | 53 |
| Median | 54 |
| Range | 20 - 80 |
| **Length of ICU stay (days)** |  |
| Mean | 18 |
| Median | 14 |
| Range | 3 - 55 |
| **Reason for admission** |  |
| Neurology | 5 (21.7%) |
| Trauma | 5 (21.7%) |
| Cardiovascular | 4 (17.4%) |
| Hepatic | 2 (8.7%) |
| Oncology | 2 (8.7%) |
| Transplant | 2 (8.7%) |
| Burns | 1 (4.3%) |
| Gastroenterology | 1 (4.3%) |
| Elective surgery | 1 (4.3%) |
| Respiratory | 1 (4.3%) |

### ICU patients were colonised by a diverse population of *Enterobacterales*

We focussed on the common commensal Gram-negative organisms *E. coli* and *Klebsiella* which can cause invasive disease (e.g. pneumonia, bloodstream infections), transmit between patients, and are known to circulate in the ICU environment (5, 6, 34). *E. coli* and *Klebsiella* have been reported to exhibit high levels of MDR, including carbapenem resistance, in other ICU environments.

WGS of organisms isolated from stool culture identified 106 *E. coli* and 17 *Klebsiella* species. Isolates that appeared to be candidate *E. coli* from colony morphology were identified as normal gut flora isolates by WGS (*Citrobacter* [n=8], *Enterobacter* [n=16]) but these were not the focus of the study. The majority of patients (n=20) were colonised with one or more Gram-negative organisms. Eighteen patients were colonised with *E. coli* during their stay. Four of these had no *E. coli* identified in their baseline sample but *E. coli* was isolated from subsequent samples during their admission. Six patients were colonised with *Klebsiella* species at some point during their stay, and of these two did not have *Klebsiella* in their baseline sample (Figure 2). Patients were frequently co-colonised with multiple species for several days (e.g., IMP2-20, IMP2-24). Three patients (IMP2-11, IMP2-15, IMP2-28) were not colonised with *E. coli* or *Klebsiella*, but were colonised with other species given a presumptive ID of *Staphylococcus, Enterococcus*, or *Pseudomonas*.

**Fig. 2.**
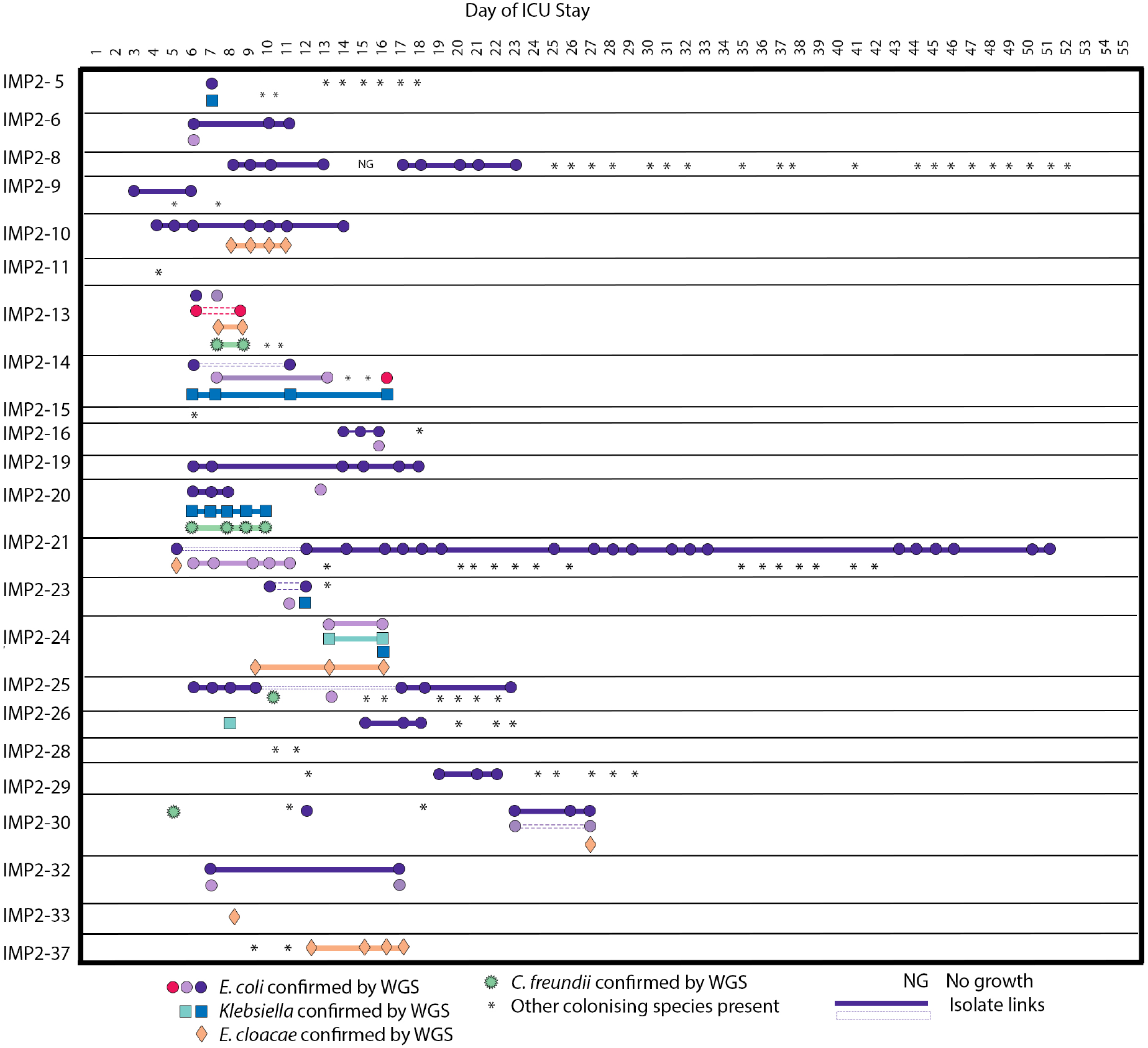
Timeline of colonising species seen during an ICU patient stay. Patients were in the ICU at different times but for illustrative purposes the timeline is the inpatient stay day number. In cases where the same ST is present multiple times in the same patient they are shown as the same colour, but these colours cannot be compared between patients. Isolate links are demonstrated with a solid line. Where this ST is interrupted by another ST, a dashed line is used. In cases where species aside from *E. coli, Klebsiella, Enterobacter* and *Citrobacter* were identified, an asterisk is used to signify these colonising isolates.

### An overview of ICU colonising *E. coli* population

WGS was used to explore the *E. coli* population on QEHB ICU, including ST diversity, ARG and plasmid carriage, and potential strain sharing between patients. Twenty-two different STs were revealed by WGS of the 106 *E. coli* isolated (Figure 3). The two most abundant STs were ST69 (n=23) and ST58 (n=22) (Tables S4 and S5). The high frequency of ST58 was as a result of repeated identification in multiple stool samples from one patient (Figure S3, Tables S4 and S5). In contrast, the abundance of ST69 was due to multiple examples of the same ST in different patients (Figure 3a). Four patients (IMP2-9, IMP2-10, IMP2-19, and IMP2-32) showed stable colonisation with the same *E. coli* strain throughout their ICU stay (Figure S6).

**Fig. 3.**
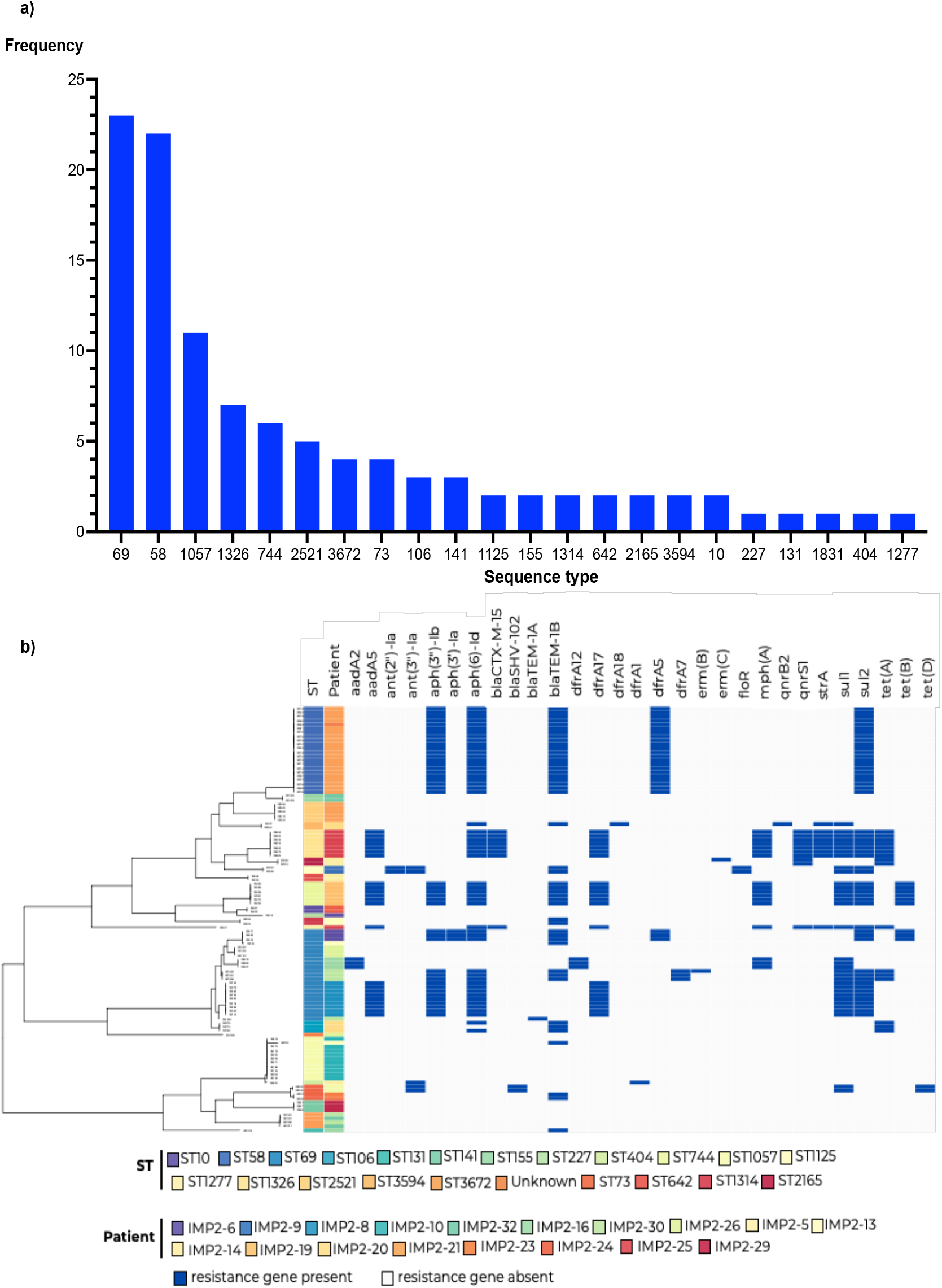
a) ST breakdown for all *E. coli* isolates (n=106) and b) the resistance gene profile for all colonising *E. coli* across all patients (n=21). Presence (navy) and absence (grey) of resistance genes are displayed alongside *E. coli* ST and patient number.

### ICU *E. coli* isolates were diverse and characterised by low levels of AMR gene carriage

Characterisation of resistance genes in this ICU *E. coli* population was carried out to ascertain whether the diversity of AMR and MDROs at QEHB ICU mirrored that reported in other ICUs worldwide. The majority of the 106 *E. coli* isolates (n=72, 68%) colonising patients were found to have one or more ARGs (Figure 3b). None of the acquired genes confer carbapenem or colistin resistance. The most common ARGs found in the colonising isolates were those that confer resistance to aminoglycosides (*aph(3’’)-Ib*/*strA* [n=43, 41%], *aph(6)-Id*/*strB* [n=54, 51%], *aad*A5 [n=23, 22%]) and sulphonamides (*sul1* and *sul2* [n=87, 82%]). In cases where there were multiple strains of the same ST (e.g. ST69), variation in ARG carriage was observed. Some, but not all, ST69 strains carried *strAB*, and there was variation between the types of *sul* and *dfr* genes carried (e.g., *sul1*/*sul2* and *dfr12*/*dfr17*). Forty-eight isolates (45%) carried *bla*_TEM_, and only ten isolates (9%) carried either of the extended-spectrum beta-lactamase (ESBL) genes *bla*_CTX-M-15_ or *bla*_SHV-102_.

The identification of resistance genes frequently found in plasmids (*bla*_CTX-M_, *bla*_TEM,_ *strAB*, various *sul* and *tet*) suggests the presence of AMR plasmids in this set of colonising ICU *E. coli* (35, 36). This inference was supported by the presence of plasmid replicons commonly found in AMR plasmids. We identified plasmid replicons belonging to families of small and large plasmids (Figure S3a). Amongst small plasmid replicons, ColRNAI types were most common, while amongst large plasmid replicons F-types (FII, FIA, FIB) dominated. As these genomes were assembled from short-read sequence data, we have not sought to further link these plasmid replicons with antibiotic resistance genes here.

The ICU *E. coli* population carried a wide range of virulence-associated genes, indicative of their potential pathogenic nature (Figure S3b). Virulence-associated genes included those for capsule, siderophores, haemolysins, P-fimbriae, and type I fimbriae.

### Low levels of AMR in other colonising species

As observed in the colonising *E. coli*, the sampled *Klebsiella* species (e.g., *Klebsiella oxytoca, Klebsiella aerogenes* and *Klebsiella pneumoniae*), were not highly drug resistant, with their most prevalent resistance genes being intrinsic resistance genes (*fosA, oqxAB*), and the intrinsic beta-lactam resistance genes, *bla*_LEN_, *bla*_OXY_ and *bla*_SHV_. All *Klebsiella* isolates sequenced carried plasmid replicons. *Klebsiella* plasmid replicons, as observed in *E. coli*, included those from small and large plasmid families (Figures S11-S13), but were less diverse. Other colonising organisms that were whole genome sequenced included *Citrobacter* and *Enterobacter* species (Figures S14-15), with a similarly low level of resistance gene carriage. *C. freundii* was the only colonising species to carry genes encoding resistance to carbapenems. Patient IMP2-20 yielded a *C. freundii* that carried *bla*_KPC-2_ from their baseline stool sample, and remained colonised by this strain for the duration of their stay. This *C. freundii* strain was not acquired by any other patients over the course of this study. The IMP2-20 colonising *C. freundii* strain carried ColRNAI, Col440I, N-type, R-type and H-type plasmid replicons along with *bla*_KPC-2_, other beta-lactamase genes, and streptomycin, sulphonamide, trimethoprim and quinolone resistance genes (Figure S14). The *bla*_KPC-2_ gene in this *C. freundii* strain was found in a 50 kb contig that also included the N-type plasmid replicon. Comparison revealed that this contig was 99.9% identical to part of the recently described plasmid pQEB1, which has been associated with QEHB since at least 2016 (37).

### ARGs with major clinical relevance were rare in ICU patient stool metagenomes

Sixteen patient stool metagenomes were screened to determine whether the sparsity of acquired ARGs in *E. coli* isolates were representative of wider gastrointestinal communities, or whether significant reservoirs of resistance were missed by examining the most abundant strains. This revealed that ARGs that confer resistance to ESBLs, carbapenems, and colistin were rare or were not present. Only two metagenomes contained *bla*_CTX-M_ genes (samples QEHB25280921 and QEH25051021), the same patient samples that produced the *bla*_CTX-M_-containing *E. coli* isolate. Intrinsic beta-lactamase genes were found in two samples: the *bla*_OXA-50_ gene of *Acinetobacter* species was in QEHB16260721, while *bla*_ADC-2_ (*Acinetobacter* species) and *bla*_OXY-1-3_ (*Klebsiella oxytoca*) were found in QEHB24170921, but without knowing the context of these it is not possible to determine the resistance phenotypes they confer. QEHB24170921 also contained *bla*_CMY-59_ and *bla*_MIR-9,_ *ampC*-type beta-lactamase genes that can confer ESBL resistance when mobilised from their original chromosomal contexts. Colistin and carbapenem resistance genes were not detected in any metagenomes, despite *bla*_KPC-2_ being found in a *C. freundii* isolated from one of the patients the metagenomes are derived from, suggesting a very low level of prevalence in that sample.

Although Gram-negative bacteria are the focus of this study, it is noteworthy that the complete set of *vanA*-type vancomycin resistance genes found in Tn*1546* were detected in sample QEHB16280721.

### Abundant *E. coli* STs were patient specific and not circulating

Circulation of STs in the QEHB ICU could facilitate the spread of AMR between vulnerable patients. To investigate the presence of circulating STs, SNP differences were calculated and used to establish potential links between strains carried by different patients. SNP analysis demonstrated that strains were associated with individual patients, with no examples found of predominant strains circulating in the ICU. This indicates that *E. coli* isolated here are most likely resident commensal strains and were not acquired by the patient during their ICU stay. The close association of strains with individual patients was demonstrated by ST69, the most abundant *E. coli* ST, where seven distinct strains of ST69 were detected in six patients (Table S4). All closely related isolates (< 13 SNPs different) were identified within the same patient, providing six individual examples of persistent colonisation with a patient-specific single strain. There was only one example of colonisation with multiple ST69 strains, in patient IMP2-30. There, the strains were notably distinct, with 2000 SNPs different. The snapshot of other species (e.g., *Klebsiella*) sequenced also demonstrated that in cases where there were multiple isolates of a single ST. In those examples, they were always isolated from the same patient but at different timepoints during the longitudinal study.

### Strain transmission on the ICU was rare

When the same ST was found in multiple patients, SNP analysis was used to determine whether strain transmission may have occurred. This SNP analysis showed strain transmission was rare in the QEHB ICU patient population. Our study highlighted a few instances of multiple patients being colonised with same ST strain, exemplified in isolates of ST58, ST1057 and ST3672 (Table S5, Figure S7).

An ST58 strain was found colonising two patients, IMP2-21 and IMP2-23. IMP2-23 arrived on the ICU after IMP2-21 and was discharged before IMP2-21. IMP2-21 arrived on the ICU colonised with ST58, whereas IMP2-23 did not have ST58 *E. coli* in their baseline stool sample and instead appeared to have acquired it during their ICU stay. Both were inpatients at the same time on the same ICU ward section, but did not have beds located near each other and never occupied the same bed location.

Two patients, IMP2-10 and IMP2-13, (Figure S7a) who stayed on the same ward but did not have overlapping stays were colonised with the same ST1057 strain (11 isolates, zero – six SNPs different). There is a possibility both patients acquired this strain whilst in hospital or in the ICU as although they had ST1057 in their baseline samples, both spent over six days on the ICU before their first stool sample, and prior to this ICU stay one patient spent 13 days on a general ward.

Patients IMP2-30 and IMP2-32, located on two different ICU wards, were colonised with the same ST3672 strain in the same month (Figure S7b). In this case there was an overlap in patient stay on the ICU (16 days). Both were admitted from the community, IMP2-32 had the ST3672 strain in their baseline sample and it was lost, being replaced by another ST3672 strain later in their stay. The ST3672 shared strain was isolated from IMP2-30 stool on day 23.

Acquisition of new *E. coli* STs appears to have occurred in eight patients during their ICU stays. Patients acquired *E. coli* from day six – 23 of their stay. Factors including time to colonisation, ward location at time of acquisition, time of ICU admission, antibiotic prescription, and length of stay all varied between these patients (Figures 4-5, S8-S10). Patients who appeared to have acquired *E. coli* STs after their initial stool sample did not all acquire strains of the same STs. Acquired STs included ST131, ST2521, ST10, ST1277, ST69, ST141, and ST3672.

**Fig. 4.**
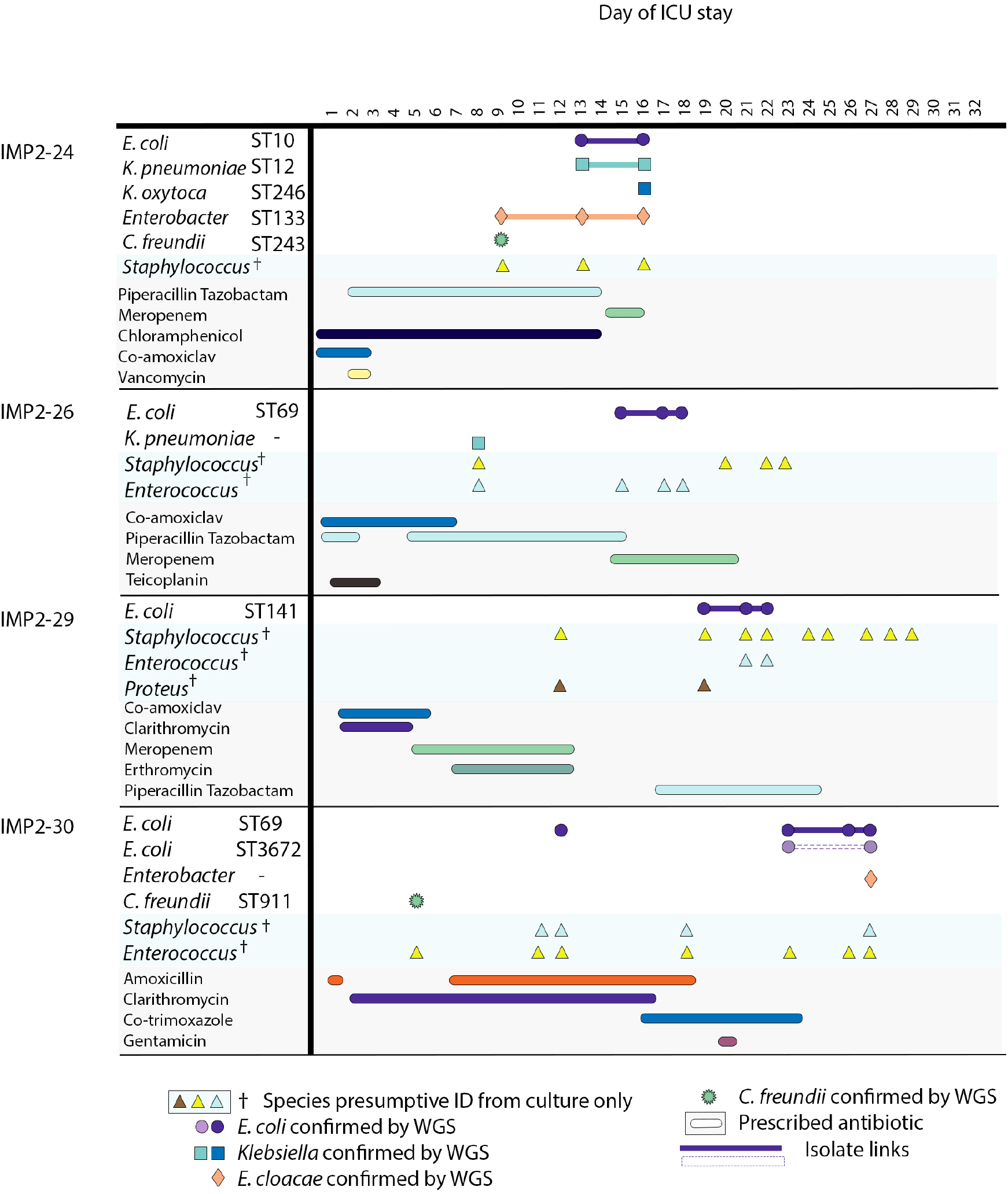
Colonising isolate timeline displaying patients who acquired *E. coli* during their stay after having no detectable *E. coli* in their baseline first stool sample. All species confirmed using WGS are displayed alongside species presumptively identified using culture plates only. Antibiotics prescribed during a patient stay is displayed below. Patients were in the ICU at differing times but for illustrative purposes the timeline is the inpatient stay day number. In cases where the same ST is present multiple times in the same patient they are shown as the same colour, but these colours cannot be compared between patients. Isolate links are demonstrated with a solid line. Where this ST is interrupted by another ST, a dashed line is used.

Patient IMP2-16 acquired the only ST131 isolate observed in this dataset (H41, Clade A). Significantly, this patient had spent time on a general ward before admission to the ICU. Their initial stool sample, produced after two weeks in the ICU, contained only *E. coli* ST69, with ST131 isolated later. Another patient, IMP2-25, transiently acquired a *bla*_CTX-M_ gene carrying ST1277 in addition to their persistent original *bla*_CTX-M-15_-carrying ST1326. The occurrence of new *E. coli* STs during the ICU stays in these eight patients is the most likely explanation of their occurrence, although time limitations of the study may have resulted in not all STs being captured.

Overall, colonisation within these patients was fluid, with strains gained and lost throughout the study (Figures 4-5, S8-10), but with minimal evidence of between-patient transmission.

## Discussion

The prevalence of MDR in ICUs globally is increasingly concerning, with multiple ICU colonisation studies identifying high incidences of MDR (5-7, 9). Here, we investigated whether the ICU at QEHB had the same burden of MDR as that observed elsewhere. Whilst limited colonisation and strain transmission was uncovered, we found no evidence of MDR in this ICU. We identified 22 different STs of colonising *E. coli* from 23 different patient samples taken from May - December 2021. Whilst this represents reasonable diversity in a single ICU, it is considerably less diverse that what has been reported elsewhere. A study in Guangzhou, China identified, for example, 59 different STs over a three-month period (6). At QEHB ICU the most abundant STs were the common commensals ST69 and ST58, which were both observed at much lower levels in the Guangzhou ICU. We investigated the colonisation dynamics of these isolates, identifying 11 patients that had been colonised by more than one *E. coli* ST during their ICU stay. There was less dynamic activity of colonising strains in QEHB ICU than what has been observed previously in nosocomial transmission events (6, 38-40). There was also no identifiable pattern in the gain (Figures 4 and S9) or loss (Figure 5) of *E. coli* strains. Sequencing an increased number of isolates per sample alongside with paired metagenomic sequencing would however have given a broader representation of colonising isolates. Overall, our data suggests that there is less transmission of isolates between patients in QEHB ICU compared to that observed in ICUs globally.

**Fig. 5.**
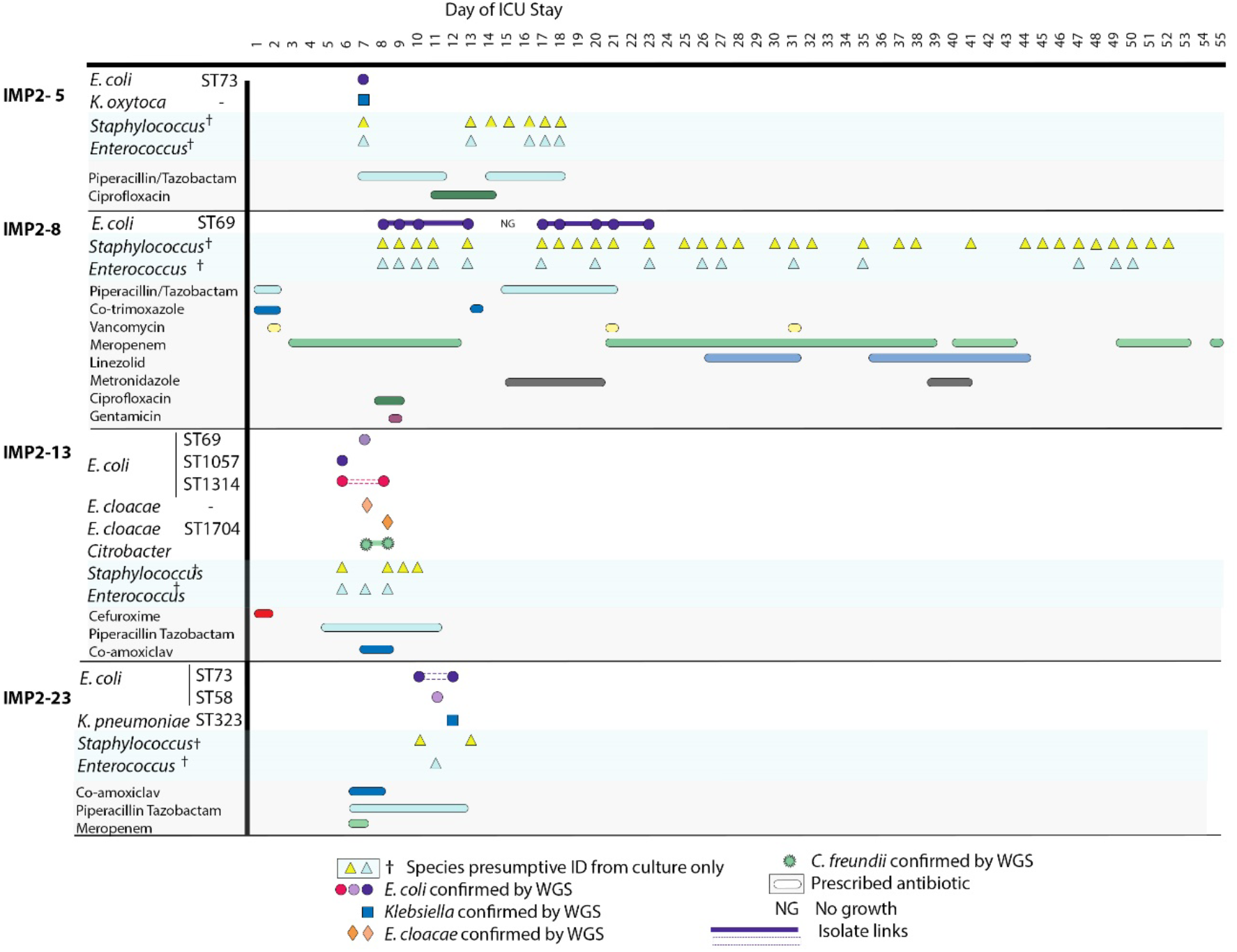
Colonising isolate timelines where *E. coli* colonisation was lost during inpatient stay. All species confirmed using WGS are displayed alongside species presumptively identified using culture plates only. Antibiotics prescribed during a patient stay is displayed below. Patients were in the ICU at differing times but for illustrative purposes the timeline is the inpatient stay day number. In cases where the same ST is present multiple times in the same patient they are shown as the same colour, but these colours cannot be compared between patients. Isolate links are demonstrated with a solid line. Where this ST is interrupted by another ST, a dashed line is used.

The lack of MDR carriage in QEHB ICU findings contrasts findings from investigations in many other countries, where multiple ICU colonisation studies have identified high rates of MDR (5-7, 9). Specifically, there was a very low level of CPE carriage. The only CPE isolated was a *bla*_KPC-2_-carrying *Citrobacter* from IMP2-20 (Figure S10), a patient who had been admitted from home. This CPE isolate was identified in the ICU baseline stool sample (day six), and it was not found in stool samples of any other patients during the study. As this *bla*_KPC-2_ gene appeared to be in a plasmid that has been strongly associated with QEHB, we cannot exclude the possibility that this *Citrobacter* strain or the plasmid it carried were acquired in the hospital over the six-day period prior to patient IMP2-20’s baseline stool sample. Similarly, very low levels of resistance to ESBLs were observed in the QEHB ICU. High occurrence of ESBLs in ICUs (including NICUs) in Europe has been shown to lead to higher mortality (49), underscoring the importance of monitoring potential outbreaks. We identified only 11 patients colonised with *E. coli* carrying β-lactamase resistance genes (e.g., *bla*_TEM,_ *bla*_SHV,_ *bla*_CTX-M_), with the majority (n=9) carrying only *bla*_TEM_. The *bla*_TEM_ gene encodes a narrow-spectrum β-lactamase which can be inhibited by β-lactamase inhibitors (e.g., clavulanic acid, tazobactam). The presence of this gene is therefore significantly less concerning than that of ESBLs as patients carrying *bla*_TEM_-carrying isolates are easily treated. Although colistin resistance is found in healthcare settings in other countries (41) there was no evidence of colistin resistance on the ICU. This observation is consistent with the low reported levels of colistin resistance in the UK more widely (15).

Within the *E. coli* species there are lineages that are particularly problematic from a healthcare perspective (40, 41), including the multidrug resistant pandemic clones ST131-H30R1 and ST131-H30Rx (42-44). Our study uncovered low numbers of STs of concern (e.g., ST69 [n=7], ST73 [n=2], ST131 [n=1]), in contrast to the higher levels found circulating in other countries (38, 42, 43). In line with reports in ICUs globally (6, 45, 46) where ST69 was detected but transmission was not frequently reported, three QEHB patients acquired ST69 on their ICU stay and no ST69 transmission was detected. The single ST131 isolated here was identified as an ST131 clade A isolate which is known to be largely drug susceptible, lacking *bla*_CTX-M_ and fluoroquinolone resistance genes (43, 47). ST131 clade A isolates are however still a prominent cause of infections in countries such as Norway (46), meaning any identification in an ICU should still be treated as a potential cause for concern. Overall, the lack of pandemic lineages here provides additional reassurance as to the low level of MDR concern in the QEHB ICU. Whilst *E. coli* was the primary focus on this study, *Klebsiella* species are also highly problematic colonisers of ICUs (7, 48-51). *Klebsiella* species can carry high levels of resistance, including plasmids encoding ESBLs (52). The identification here of a colonising ST20 *K. pneumoniae* is consistent with previous reports where ST20 *K. pneumoniae* high risk clones caused nosocomial outbreaks (52, 53). The strain isolated however carry any of the AMR genes of concern (e.g. *bla*_CTX_ and *bla*_NDM_) that have been found in other studies (54). Whilst this strain is therefore non-MDR, it highlights the importance of routine surveillance to monitor the potential gain of MDR plasmids by high-risk clones in a hospital setting. WGS is a critical tool for this. It was employed here throughout an inpatient stay to obtain and characterise strain-level colonisation dynamics, and it is critical for detecting and understanding the presence of circulating strains. WGS in combination with longitudinal sequential sampling, as employed here, can also be used to assess the quality of infection prevention control precautions.

This study was carried out when the UK and QEHB were under COVID-19 restrictions, representing a snapshot of colonisation in unusual, highly controlled conditions. Patients were for example shielded from potential colonisation opportunities as travel and personal interactions were limited prior to admission. Visitors were limited, as was patient mobility within the hospital, and extensive personal protective equipment was worn. This controlled patient exposure was in complete contrast to the freedom of movement seen in previous hospital studies (15), and our results should be considered in this context when compared to previous colonisation studies in QEHB (15) and elsewhere in the UK (38). There was also a general reduction in global travel, reducing the likelihood of colonisation by ESBL-*E. coli* or CPEs. Isolates detected in the first, baseline stool sample are also more likely to reflect local community carriage at the time of admission as opposed to a more national picture that might have been seen previously.

The overview of specific Gram-negative organisms, including *E. coli* and *Klebsiella*, detailed here gives a better understanding of the wider picture of colonisation dynamics in a large ICU ward during a period of COVID-19 restrictions. Tracking and characterising the AMR carriage found in colonising *E. coli* uncovered low levels of resistance, underscoring the importance of robust infection control measures.

## Supporting information

Supplementary isolate list

## Abbreviations

MDR: Multidrug resistance
MDRO: Multidrug resistant organism
QEHB: Queen Elizabeth Hospital Birmingham
ICU: intensive care unit
IPC: Infection Prevention and Control
UHB: University Hospitals Birmingham, NHS Foundation Trust
ST: Sequence type
WGS: Whole genome sequencing
ARG: Antibiotic resistance gene
CPE: Carbapenemase-producing Enterobacterales

## Author contributions

AM, TW, WVS: Conceptualisation

AES, AC, LR: Data curation

TW, AC, LR: Project administration

AES, RJH, RAM: formal analysis

AES: Visualisation

AES, RAM, RJH, AM: Writing – original draft

All authors: Writing – review & editing

## Ethical approval

The trial was approved by the Yorkshire & The Humber - Leeds East Research Ethics Committee (Reference, 20/YH/0067).

## Acknowledgements

The authors wish to acknowledge the support of the following research nurses and staff at Queen Elizabeth Hospital, University Hospitals Birmingham NHS Trust for the help with sample collection for this study: Laura Mee, Hazel Smith, Natalie Dooley, Emma Fellows, Amy Clark, Stephanie Goundry, Christopher Sheridan, Puja Mulholland, Colin Bergin, Elaine Spruce, Saffron King, Alex Newton-Cox, Samantha Harkett, Connor Bentley, Aoife Neal, Tyreeck Joseph, and Sophia Beddows.

