## Supplementary figures and tables for "Longitudinal genomic surveillance of a UK intensive care unit shows a lack of patient colonisation by multi-drug resistant Gram-negative pathogens"

**Table S1.** IMPACT2 patient information.

| Sample Number | Patient Number | Gender | Age | Length of ICU Stay (days) | Overarching admission category | Admission from |
|---|---|---|---|---|---|---|
| QEHB05 | IMP2-05 | F | 45 | 19 | Neurology | Other hospital |
| QEHB06 | IMP2-06 | F | 29 | 11 | Hepatic | Other hospital |
| QEHB08 | IMP2-08 | F | 47 | 55 | Transplant | Community |
| QEHB09 | IMP2-09 | M | 23 | 5 | Burns | Community |
| QEHB10 | IMP2-10 | M | 41 | 13 | Cardiovascular | Community |
| QEHB11 | IMP2-11 | M | 77 | 3 | Cardiovascular | 4 days on QEHB general ward |
| QEHB13 | IMP2-13 | M | 50 | 10 | Neurology | 10 days on QEHB general ward |
| QEHB14 | IMP2-14 | F | 57 | 14 | Neurology | Other hospital |
| QEHB15 | IMP2-15 | M | 79 | 5 | Cardiovascular | Community |
| QEHB16 | IMP2-16 | M | 80 | 7 | Neurology | 10 days on QEHB general ward |
| QEHB19 | IMP2-19 | M | 68 | 20 | Trauma | Community |
| QEHB20 | IMP2-20 | M | 73 | 14 | Oncology | Community |
| QEHB21 | IMP2-21 | F | 74 | 51 | Trauma | Community |
| QEHB23 | IMP2-23 | F | 53 | 14 | Cardiovascular | Community |
| QEHB24 | IMP2-24 | M | 20 | 16 | Trauma | Community |
| QEHB25 | IMP2-25 | F | 61 | 24 | Oncology | 1 day on QEHB general ward |
| QEHB26 | IMP2-26 | M | 51 | 22 | Gastroenterology | Other hospital |
| QEHB28 | IMP2-28 | F | 54 | 9 | Elective surgery | Community |
| QEHB29 | IMP2-29 | M | 51 | 30 | Neurology | Community |
| QEHB30 | IMP2-30 | M | 55 | 27 | Respiratory | Community |
| QEHB32 | IMP2-32 | M | 56 | 17 | Trauma | Community |
| QEHB33 | IMP2-33 | M | 25 | 8 | Trauma | Community |
| QEHB37 | IMP2-37 | M | 44 | 16 | Transplant | Community |

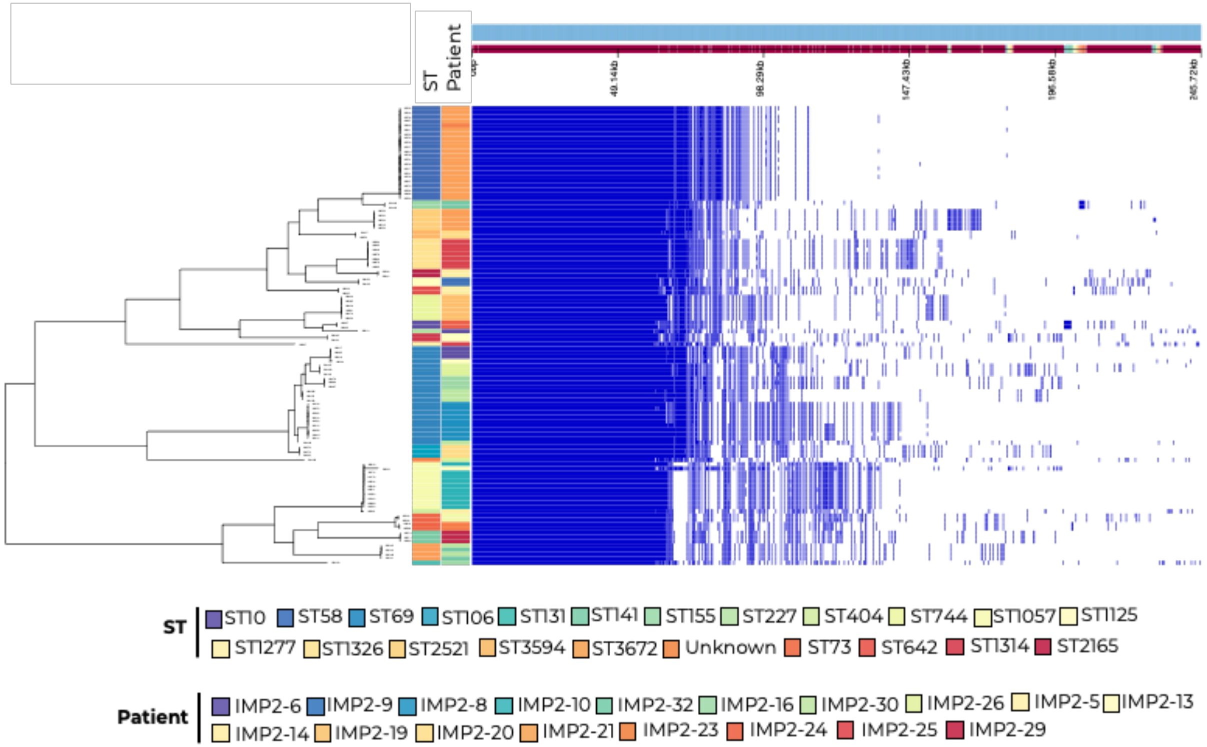


**Fig. S2.** *E. coli* core gene alignment phylogeny with Panaroo core gene presence absence profile in navy displayed alongside patient number and *E. coli* ST.

a)

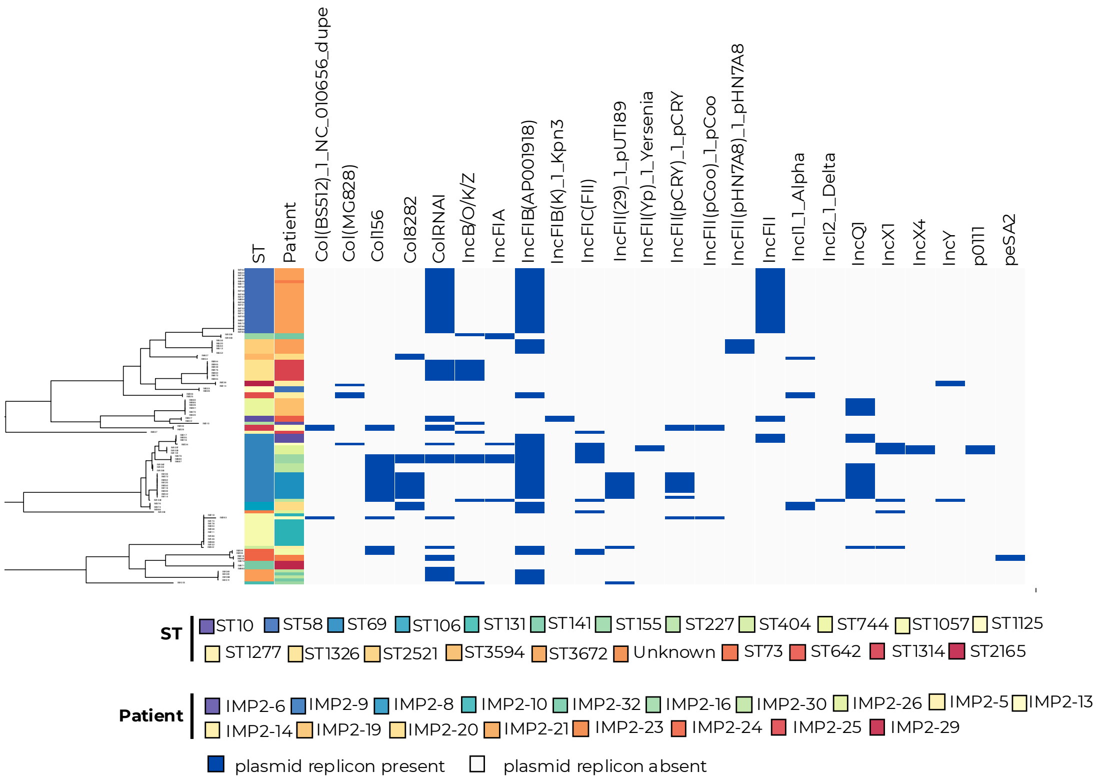
ST
Patient
Col(BS512)_1_NC_010656_dupe
Col(MG828)
Col156
Col8282
ColRNAI
IncB/O/K/Z
IncFIA
IncFIB(AP001918)
IncFIB(K)_1_Kpn3
IncFIC(FII)
IncFII(29)_1_pUTI89
IncFII(Yp)_1_Yersenia
IncFII(pCRY)_1_pCRY
IncFII(pCoo)_1_pCoo
IncFII(pHN7A8)_1_pHN7A8
IncFII
IncI1_1_Alpha
IncI2_1_Delta
IncQ1
IncX1
IncX4
IncY
p0111
peSA2
ST
ST10 ST58 ST69 ST106 ST131 ST141 ST155 ST227 ST404 ST744 ST1057 ST1125
ST1277 ST1326 ST2521 ST3594 ST3672 Unknown ST73 ST642 ST1314 ST2165
Patient
IMP2-6 IMP2-9 IMP2-8 IMP2-10 IMP2-32 IMP2-16 IMP2-30 IMP2-26 IMP2-5 IMP2-13
IMP2-14 IMP2-19 IMP2-20 IMP2-21 IMP2-23 IMP2-24 IMP2-25 IMP2-29
plasmid replicon present
plasmid replicon absent

b)

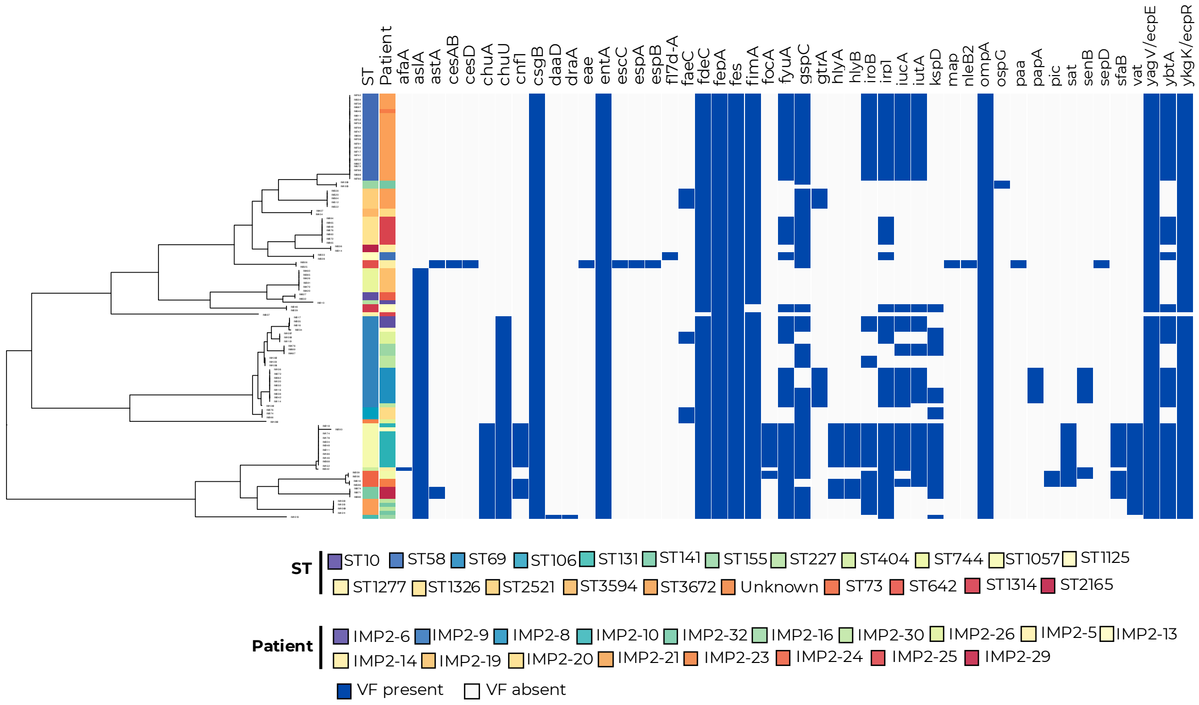


**Fig. S3.** A) Plasmid replicon profile and b) virulence factor profile for all colonising *E. coli* displayed alongside ST and patient number. Plasmid replicon/virulence gene presence (navy) and absence (grey) are indicated. A selection of virulence genes are displayed here, representing those normally seen in pathogenic *E. coli*.

**Table S4** SNP difference matrix of ST69 strains*

| Isolate | Patient | IM476 | IM487 | IM889 | IM234 | IM1093 | IM1097 | IM1101 | IM1049 | IM1051 | IM1056 | IM1062 | IM005 | IM016 | IM017 | IM039 | IM042 | IM050 | IM062 | IM072 | IM106 | IM114 | IM116 | IM120 |
|---|---|---|---|---|---|---|---|---|---|---|---|---|---|---|---|---|---|---|---|---|---|---|---|---|
| IM476 | IMP2-16 | 0 | 0 | 0 | 7651 | 6049 | 6050 | 6045 | 6766 | 4701 | 4700 | 4701 | 7563 | 7566 | 7563 | 8095 | 8093 | 8096 | 8096 | 8095 | 8097 | 8097 | 8096 | 8095 |
| IM487 | IMP2-16 | 0 | 0 | 0 | 7651 | 6049 | 6050 | 6045 | 6766 | 4701 | 4700 | 4701 | 7563 | 7566 | 7563 | 8095 | 8093 | 8096 | 8096 | 8095 | 8097 | 8097 | 8096 | 8095 |
| IM889 | IMP2-16 | 0 | 0 | 0 | 7651 | 6049 | 6050 | 6045 | 6766 | 4701 | 4700 | 4701 | 7563 | 7566 | 7563 | 8095 | 8093 | 8096 | 8096 | 8095 | 8097 | 8097 | 8096 | 8095 |
| IM234 | IMP2-13 | 7651 | 7651 | 7651 | 0 | 6566 | 6567 | 6562 | 8201 | 6044 | 6043 | 6044 | 406 | 409 | 406 | 7834 | 7832 | 7835 | 7835 | 7834 | 7836 | 7836 | 7835 | 7834 |
| IM1093 | IMP2-26 | 6049 | 6049 | 6049 | 6566 | 0 | 13 | 4 | 6076 | 3957 | 3956 | 3957 | 6299 | 6302 | 6299 | 6370 | 6368 | 6371 | 6371 | 6370 | 6372 | 6372 | 6371 | 6370 |
| IM1097 | IMP2-26 | 6050 | 6050 | 6050 | 6567 | 13 | 0 | 9 | 6077 | 3958 | 3957 | 3958 | 6300 | 6303 | 6300 | 6371 | 6369 | 6372 | 6372 | 6371 | 6373 | 6373 | 6372 | 6371 |
| IM1101 | IMP2-26 | 6045 | 6045 | 6045 | 6562 | 4 | 9 | 0 | 6072 | 3953 | 3952 | 3953 | 6295 | 6298 | 6295 | 6366 | 6364 | 6367 | 6367 | 6366 | 6368 | 6368 | 6367 | 6366 |
| IM1049 | IMP2-30 | 6766 | 6766 | 6766 | 8201 | 6076 | 6077 | 6072 | 0 | 2363 | 2362 | 2363 | 7929 | 7932 | 7929 | 1691 | 1689 | 1692 | 1692 | 1691 | 1693 | 1698 | 1692 | 1693 |
| IM1051 | IMP2-30 | 4701 | 4701 | 4701 | 6044 | 3957 | 3958 | 3953 | 2363 | 0 | 1 | 0 | 5798 | 5801 | 5798 | 3782 | 3780 | 3783 | 3783 | 3782 | 3784 | 3784 | 3783 | 3782 |
| IM1056 | IMP2-30 | 4700 | 4700 | 4700 | 6043 | 3956 | 3957 | 3952 | 2362 | 1 | 0 | 1 | 5797 | 5800 | 5797 | 3781 | 3779 | 3782 | 3782 | 3781 | 3783 | 3783 | 3782 | 3781 |
| IM1062 | IMP2-30 | 4701 | 4701 | 4701 | 6044 | 3957 | 3958 | 3953 | 2363 | 0 | 1 | 0 | 5798 | 5801 | 5798 | 3782 | 3780 | 3783 | 3783 | 3782 | 3784 | 3784 | 3783 | 3782 |
| IM005 | IMP2-6 | 7563 | 7563 | 7563 | 406 | 6299 | 6300 | 6295 | 7929 | 5798 | 5797 | 5798 | 0 | 3 | 0 | 7581 | 7579 | 7582 | 7582 | 7581 | 7583 | 7583 | 7582 | 7581 |
| IM016 | IMP2-6 | 7566 | 7566 | 7566 | 409 | 6302 | 6303 | 6298 | 7932 | 5801 | 5800 | 5801 | 3 | 0 | 3 | 7584 | 7582 | 7585 | 7585 | 7584 | 7586 | 7586 | 7585 | 7584 |
| IM017 | IMP2-6 | 7563 | 7563 | 7563 | 406 | 6299 | 6300 | 6295 | 7929 | 5798 | 5797 | 5798 | 0 | 3 | 0 | 7581 | 7579 | 7582 | 7582 | 7581 | 7583 | 7583 | 7582 | 7581 |
| IM039 | IMP2-8 | 8095 | 8095 | 8095 | 7834 | 6370 | 6371 | 6366 | 1691 | 3782 | 3781 | 3782 | 7581 | 7584 | 7581 | 0 | 4 | 3 | 3 | 4 | 4 | 9 | 3 | 4 |
| IM042 | IMP2-8 | 8093 | 8093 | 8093 | 7832 | 6368 | 6369 | 6364 | 1689 | 3780 | 3779 | 3780 | 7579 | 7582 | 7579 | 4 | 0 | 5 | 5 | 6 | 6 | 11 | 5 | 6 |
| IM050 | IMP2-8 | 8096 | 8096 | 8096 | 7835 | 6371 | 6372 | 6367 | 1692 | 3783 | 3782 | 3783 | 7582 | 7585 | 7582 | 3 | 5 | 0 | 0 | 1 | 1 | 8 | 4 | 1 |
| IM062 | IMP2-8 | 8096 | 8096 | 8096 | 7835 | 6371 | 6372 | 6367 | 1692 | 3783 | 3782 | 3783 | 7582 | 7585 | 7582 | 3 | 5 | 0 | 0 | 1 | 1 | 8 | 4 | 1 |
| IM072 | IMP2-8 | 8095 | 8095 | 8095 | 7834 | 6370 | 6371 | 6366 | 1691 | 3782 | 3781 | 3782 | 7581 | 7584 | 7581 | 4 | 6 | 1 | 1 | 0 | 2 | 9 | 5 | 2 |
| IM106 | IMP2-8 | 8097 | 8097 | 8097 | 7836 | 6372 | 6373 | 6368 | 1693 | 3784 | 3783 | 3784 | 7583 | 7586 | 7583 | 4 | 6 | 1 | 1 | 2 | 0 | 9 | 5 | 2 |
| IM114 | IMP2-8 | 8097 | 8097 | 8097 | 7836 | 6372 | 6373 | 6368 | 1698 | 3784 | 3783 | 3784 | 7583 | 7586 | 7583 | 9 | 11 | 8 | 8 | 9 | 9 | 0 | 10 | 9 |
| IM116 | IMP2-8 | 8096 | 8096 | 8096 | 7835 | 6371 | 6372 | 6367 | 1692 | 3783 | 3782 | 3783 | 7582 | 7585 | 7582 | 3 | 5 | 4 | 4 | 5 | 5 | 10 | 0 | 5 |
| IM120 | IMP2-8 | 8095 | 8095 | 8095 | 7834 | 6370 | 6371 | 6366 | 1693 | 3782 | 3781 | 3782 | 7581 | 7584 | 7581 | 4 | 6 | 1 | 1 | 2 | 2 | 9 | 5 | 0 |

*Isolates with less than 13 SNPs different are highlighted in red. Patient number is displayed alongside SNP differences. White and grey banding is used to display different patients by rows. Only IMP2-6 and IMP2-8 had overlapping ICU stays.

**Table S5** SNP difference matrix of ST58 strains*

| Isolate | Patient | IM504 | IM572 | IM646 | IM688 | IM694 | IM708 | IM717 | IM732 | IM736 | IM741 | IM747 | IM752 | IM759 | IM762 | IM780 | IM786 | IM791 | IM799 | IM807 | IM811 | IM687 | IM700 |
|---|---|---|---|---|---|---|---|---|---|---|---|---|---|---|---|---|---|---|---|---|---|---|---|
| IM504 | IMP2-21 | 0 | 0 | 0 | 0 | 0 | 0 | 0 | 0 | 0 | 0 | 0 | 0 | 0 | 0 | 0 | 0 | 0 | 0 | 0 | 0 | 1 | 1 |
| IM572 | IMP2-21 | 0 | 0 | 0 | 0 | 0 | 0 | 0 | 0 | 0 | 0 | 0 | 0 | 0 | 0 | 0 | 0 | 0 | 0 | 0 | 0 | 1 | 1 |
| IM708 | IMP2-21 | 0 | 0 | 0 | 0 | 0 | 0 | 0 | 0 | 0 | 0 | 0 | 0 | 0 | 0 | 0 | 0 | 0 | 0 | 0 | 0 | 1 | 1 |
| IM688 | IMP2-21 | 0 | 0 | 0 | 0 | 0 | 0 | 0 | 0 | 0 | 0 | 0 | 0 | 0 | 0 | 0 | 0 | 0 | 0 | 0 | 0 | 1 | 1 |
| IM694 | IMP2-21 | 0 | 0 | 0 | 0 | 0 | 0 | 0 | 0 | 0 | 0 | 0 | 0 | 0 | 0 | 0 | 0 | 0 | 0 | 0 | 0 | 1 | 1 |
| IM717 | IMP2-21 | 0 | 0 | 0 | 0 | 0 | 0 | 0 | 0 | 0 | 0 | 0 | 0 | 0 | 0 | 0 | 0 | 0 | 0 | 0 | 0 | 1 | 1 |
| IM732 | IMP2-21 | 0 | 0 | 0 | 0 | 0 | 0 | 0 | 0 | 0 | 0 | 0 | 0 | 0 | 0 | 0 | 0 | 0 | 0 | 0 | 0 | 1 | 1 |
| IM736 | IMP2-21 | 0 | 0 | 0 | 0 | 0 | 0 | 0 | 0 | 0 | 0 | 0 | 0 | 0 | 0 | 0 | 0 | 0 | 0 | 0 | 0 | 1 | 1 |
| IM741 | IMP2-21 | 0 | 0 | 0 | 0 | 0 | 0 | 0 | 0 | 0 | 0 | 0 | 0 | 0 | 0 | 0 | 0 | 0 | 0 | 0 | 0 | 1 | 1 |
| IM646 | IMP2-23 | 0 | 0 | 0 | 0 | 0 | 0 | 0 | 0 | 0 | 0 | 0 | 0 | 0 | 0 | 0 | 0 | 0 | 0 | 0 | 0 | 1 | 1 |
| IM747 | IMP2-21 | 0 | 0 | 0 | 0 | 0 | 0 | 0 | 0 | 0 | 0 | 0 | 0 | 0 | 0 | 0 | 0 | 0 | 0 | 0 | 0 | 1 | 1 |
| IM752 | IMP2-21 | 0 | 0 | 0 | 0 | 0 | 0 | 0 | 0 | 0 | 0 | 0 | 0 | 0 | 0 | 0 | 0 | 0 | 0 | 0 | 0 | 1 | 1 |
| IM759 | IMP2-21 | 0 | 0 | 0 | 0 | 0 | 0 | 0 | 0 | 0 | 0 | 0 | 0 | 0 | 0 | 0 | 0 | 0 | 0 | 0 | 0 | 1 | 1 |
| IM762 | IMP2-21 | 0 | 0 | 0 | 0 | 0 | 0 | 0 | 0 | 0 | 0 | 0 | 0 | 0 | 0 | 0 | 0 | 0 | 0 | 0 | 0 | 1 | 1 |
| IM780 | IMP2-21 | 0 | 0 | 0 | 0 | 0 | 0 | 0 | 0 | 0 | 0 | 0 | 0 | 0 | 0 | 0 | 0 | 0 | 0 | 0 | 0 | 1 | 1 |
| IM786 | IMP2-21 | 0 | 0 | 0 | 0 | 0 | 0 | 0 | 0 | 0 | 0 | 0 | 0 | 0 | 0 | 0 | 0 | 0 | 0 | 0 | 0 | 1 | 1 |
| IM791 | IMP2-21 | 0 | 0 | 0 | 0 | 0 | 0 | 0 | 0 | 0 | 0 | 0 | 0 | 0 | 0 | 0 | 0 | 0 | 0 | 0 | 0 | 1 | 1 |
| IM799 | IMP2-21 | 0 | 0 | 0 | 0 | 0 | 0 | 0 | 0 | 0 | 0 | 0 | 0 | 0 | 0 | 0 | 0 | 0 | 0 | 0 | 0 | 1 | 1 |
| IM807 | IMP2-21 | 0 | 0 | 0 | 0 | 0 | 0 | 0 | 0 | 0 | 0 | 0 | 0 | 0 | 0 | 0 | 0 | 0 | 0 | 0 | 0 | 1 | 1 |
| IM811 | IMP2-21 | 0 | 0 | 0 | 0 | 0 | 0 | 0 | 0 | 0 | 0 | 0 | 0 | 0 | 0 | 0 | 0 | 0 | 0 | 0 | 0 | 1 | 1 |
| IM687 | IMP2-21 | 1 | 1 | 1 | 1 | 1 | 1 | 1 | 1 | 1 | 1 | 1 | 1 | 1 | 1 | 1 | 1 | 1 | 1 | 1 | 1 | 0 | 2 |
| IM700 | IMP2-21 | 1 | 1 | 1 | 1 | 1 | 1 | 1 | 1 | 1 | 1 | 1 | 1 | 1 | 1 | 1 | 1 | 1 | 1 | 1 | 1 | 2 | 0 |

*Isolates with 0 SNPs different are highlighted in yellow. Patient number is displayed alongside SNP differences. White and grey banding is used to display different patients by rows. IM646 was isolated a day after IM741.

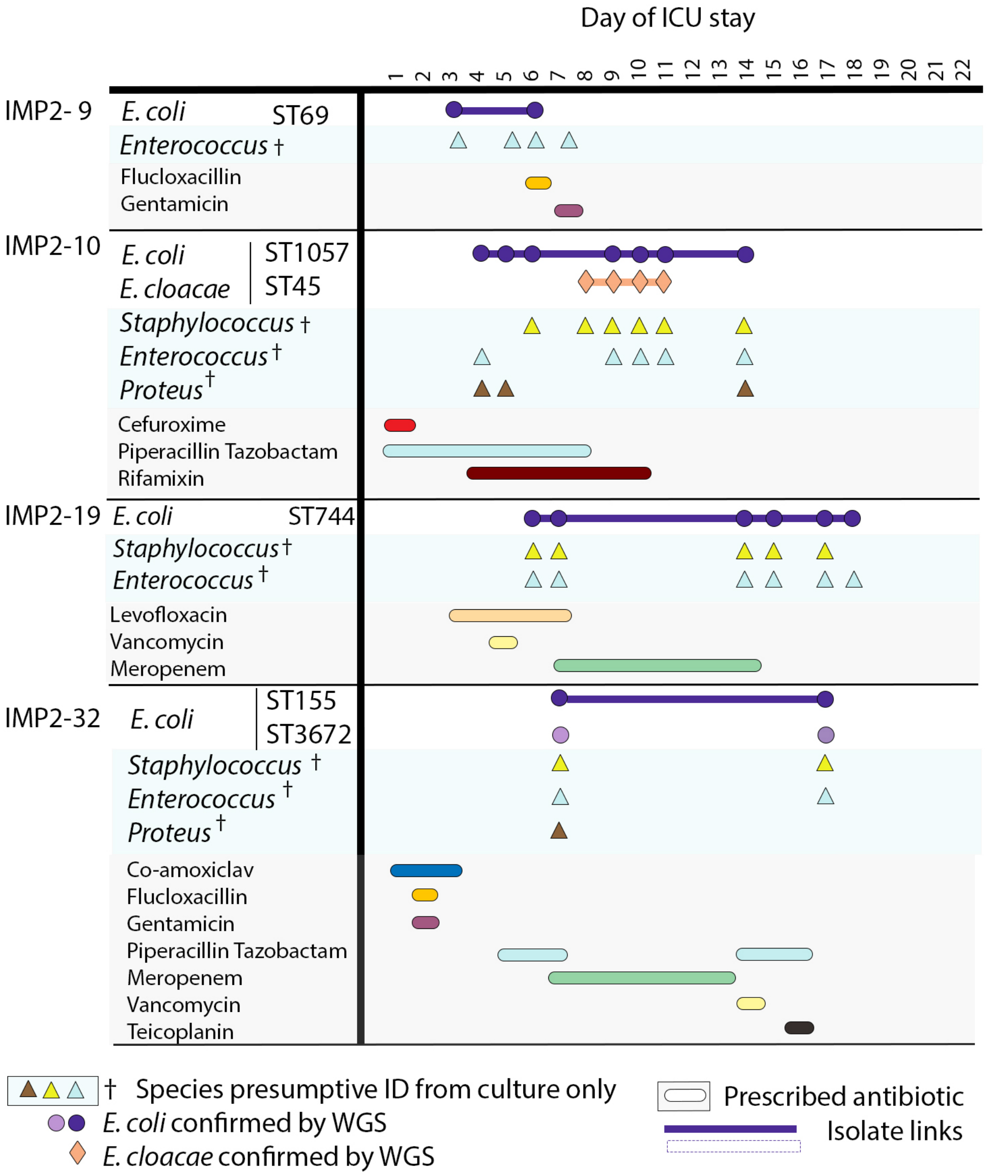


**Fig. S6** Colonising isolates in patients where *E. coli* colonisation was stable during inpatient stay. All species confirmed using WGS are displayed alongside species presumptively identified using culture plates only. Antibiotics prescribed during a patient stay is displayed below. Patients were in the ICU at differing times but for the purpose of this figure the timeline is the inpatient stay day number. In cases where STs are the same they are shown on this figure as the same colour when they occurred in the same patient, but these colours cannot be compared across patients. Isolate links demonstrated with a solid line where colonisation occurs. A dashed line is used where this ST strain is interrupted by another ST.

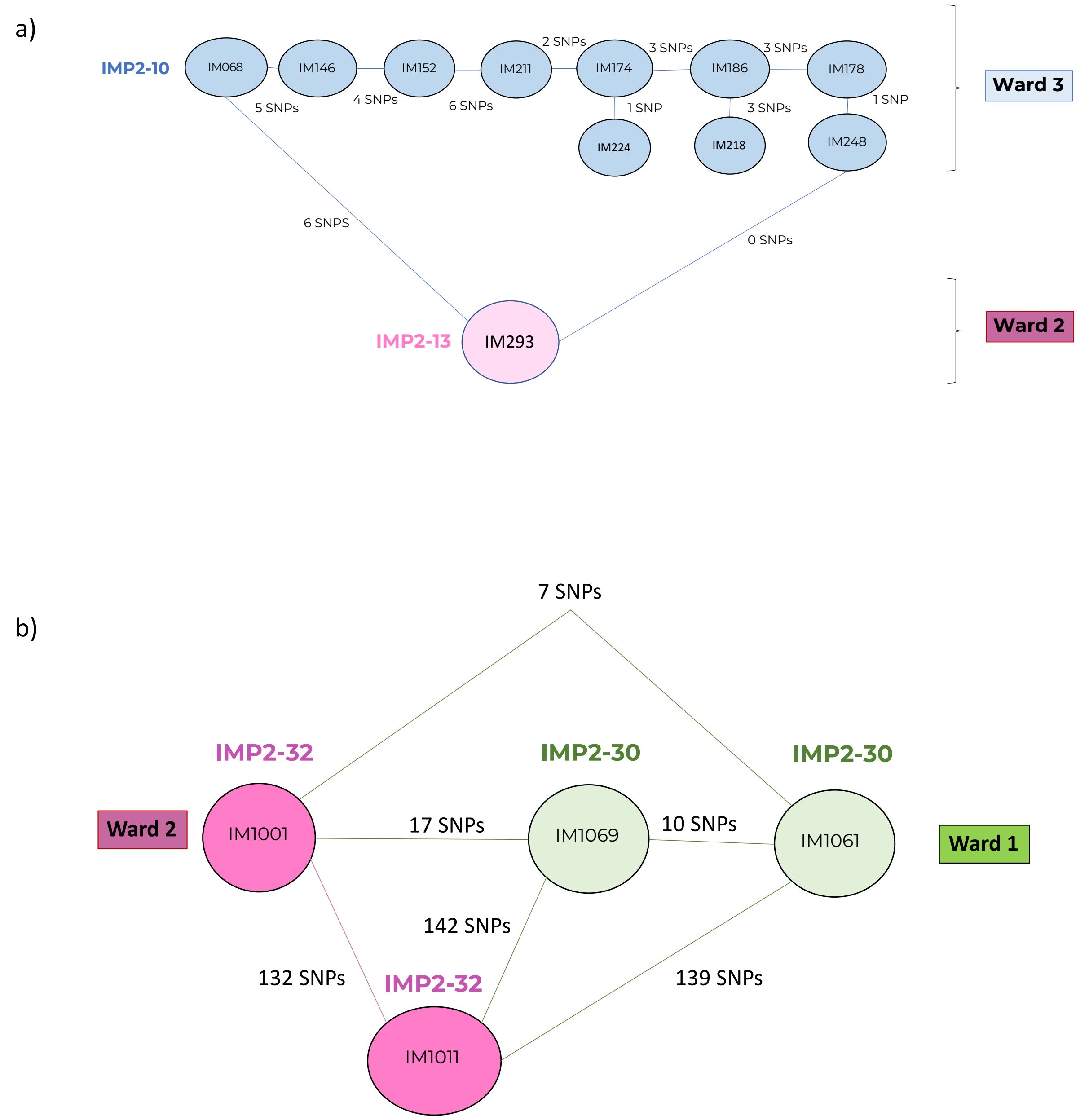


**Fig. S7** - Colonisation of different patients with the same *E. coli* strain of a) ST1057 and b) ST3672. IM number refers to isolate number and IMP2 number is patient number. SNP differences and ward location (Ward 1 /Ward 2 /Ward 3) are displayed. Ward name where the patient was staying at the time of isolation is linked to the colour of isolate from the ward (Ward 1: Green, Ward 2: Pink, Ward 3: Blue). IM293 was isolated 29 days after IM068. IM1001 and IM1011 were isolated 23 and 33 days after IM1069 respectively.

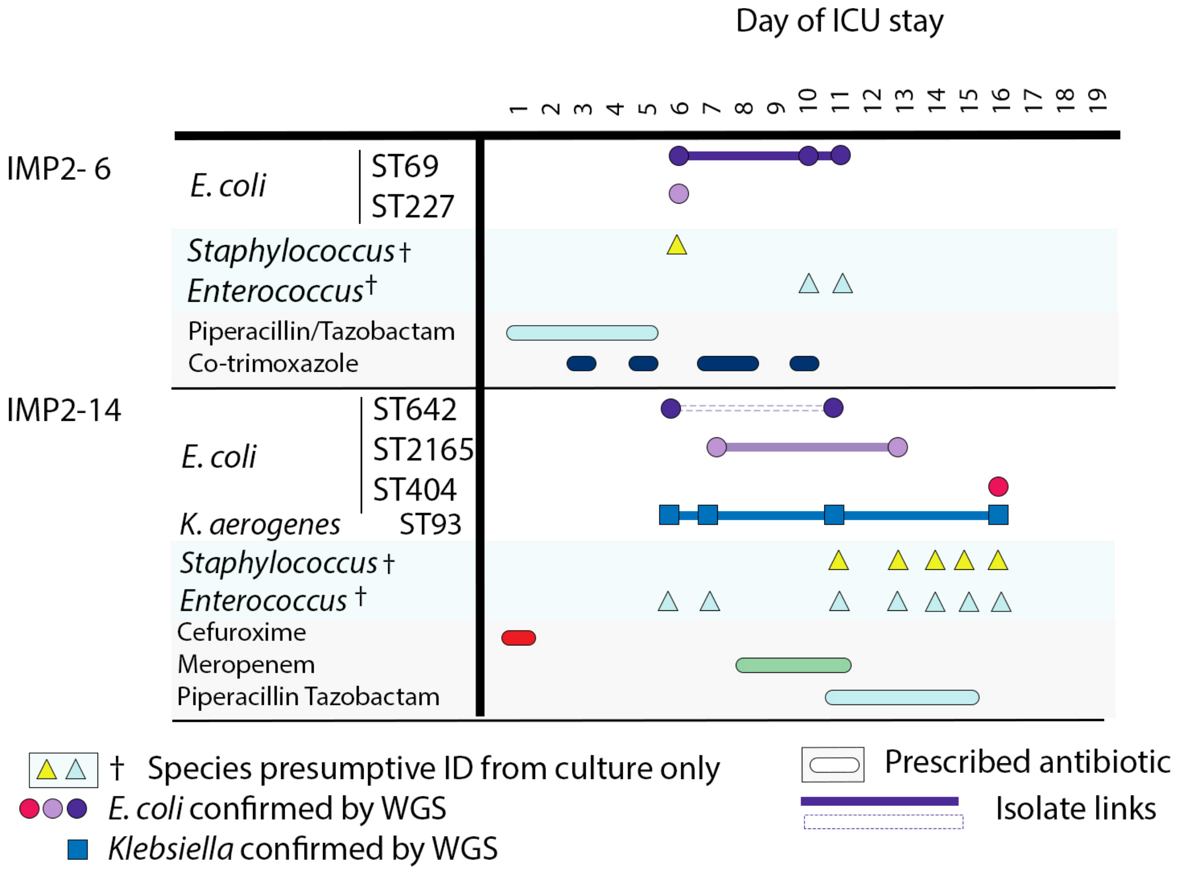


**Fig. S8** Fluctuating colonisation in ICU patients. All species confirmed using WGS are displayed alongside species presumptively identified using culture plates only. Antibiotics prescribed during a patient stay is displayed below. Patients were in the ICU at differing times but for the purpose of this figure the timeline is the inpatient stay day number. In cases where STs are the same they are shown on this figure as the same colour when they occurred in the same patient, but these colours cannot be compared across patients. Isolate links demonstrate with a solid line where colonisation occurs with a solid line and where this ST strain is interrupted by another ST, a dashed line is used.

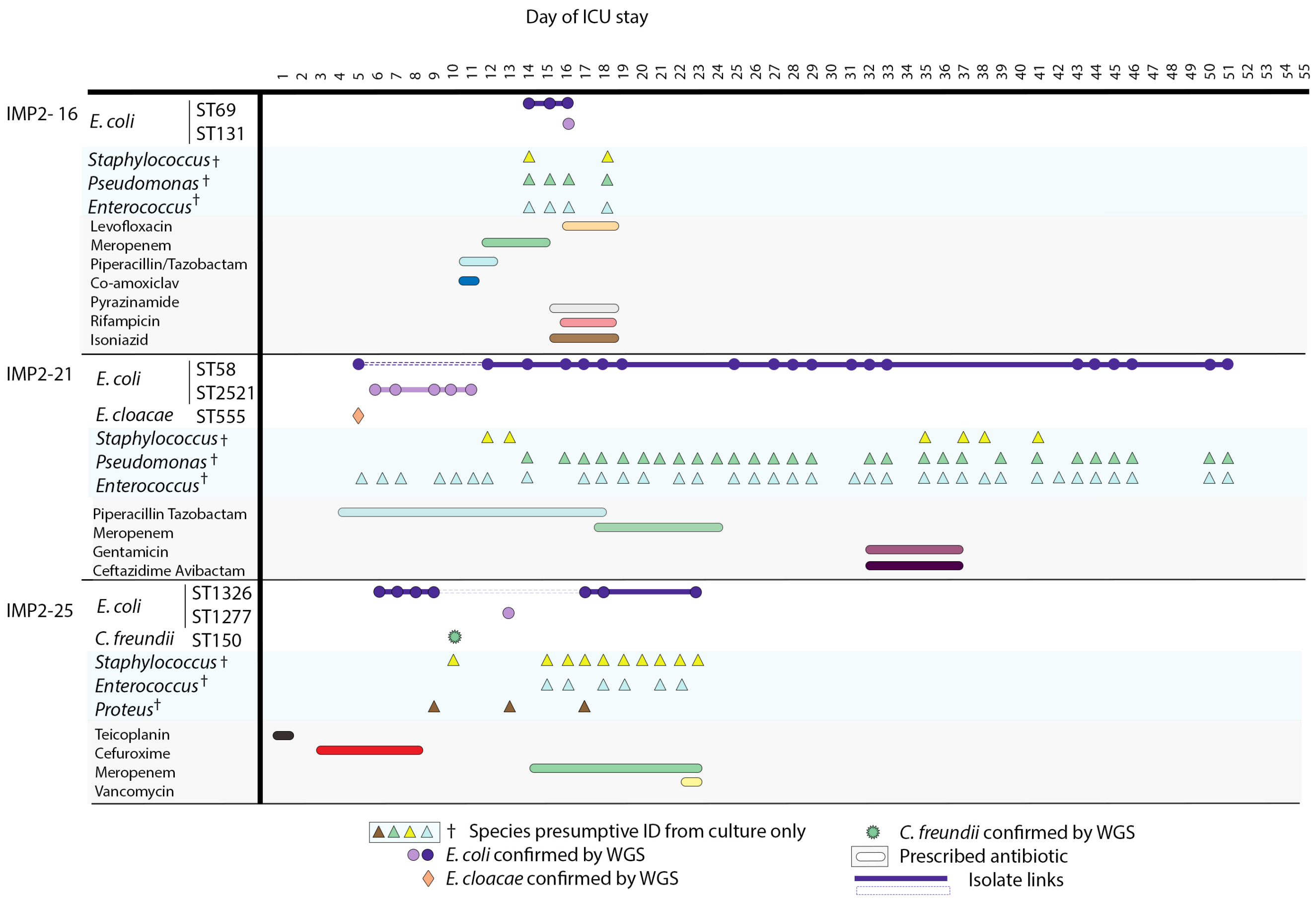


**Fig. S9** Colonising isolate timeline where the patient acquired a new *E. coli* ST during their stay. All species confirmed using WGS are displayed alongside species presumptively identified using culture plates only. Antibiotics prescribed during a patient stay is displayed below. Patients were in the ICU at differing times but for the purpose of this figure the timeline is the inpatient stay day number. In cases where STs are the same they are shown on this figure as the same colour when they occurred in the same patient, but these colours cannot be compared across patients. Isolate links demonstrate with a solid line where colonisation occurs with a solid line and where this ST strain is interrupted by another ST, a dashed line is used.

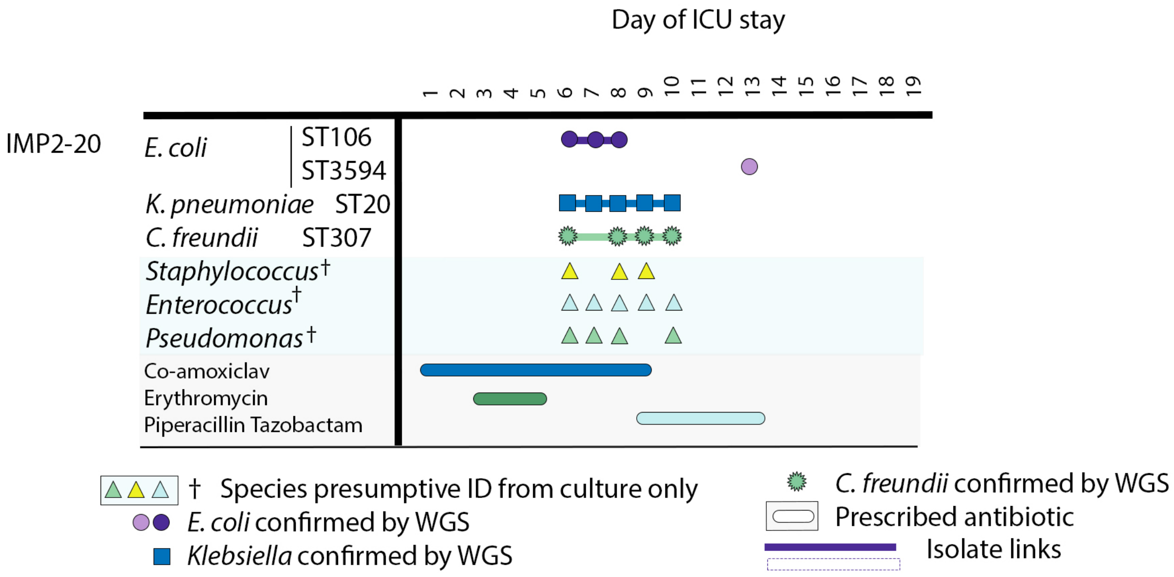


**Fig. S10** IMP2-20 timeline where an acquired *E. coli* appeared to overgrow other initial colonising species. All species confirmed using WGS are displayed alongside species presumptively identified using culture plates only. Antibiotics prescribed during a patient stay is displayed below. Patients were in the ICU at differing times but for the purpose of this figure the timeline is the inpatient stay day number. In cases where STs are the same they are shown on this figure as the same colour when they occurred in the same patient, but these colours cannot be compared across patients. Isolate links demonstrate with a solid line where colonisation occurs with a solid line and where this ST strain is interrupted by another ST, a dashed line is used.

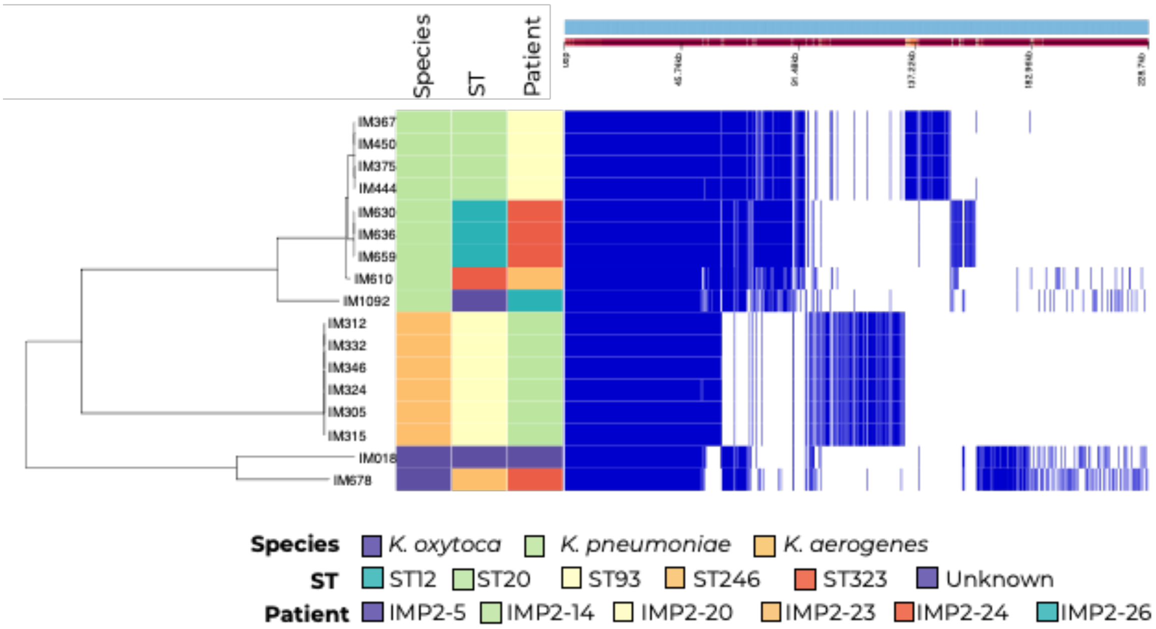


**Fig. S11** *Klebsiella* core gene alignment phylogeny with Panaroo core gene presence absence profile, patient number and *E. coli* ST displayed alongside. Blue is core gene presence and white is core gene absence. Phylogenetic tree is midpoint rooted with increasing nodes.

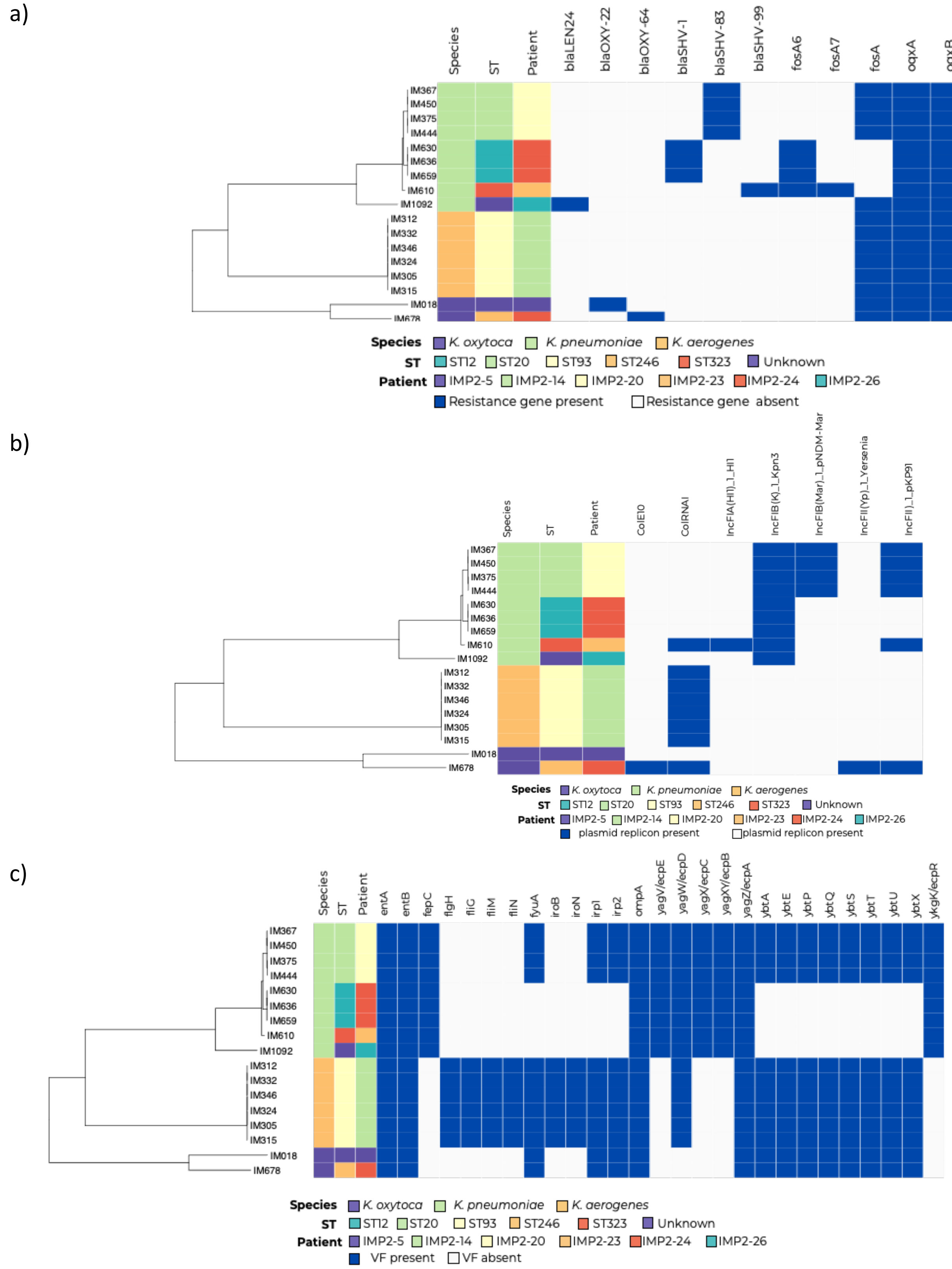


**Fig. S12** a) Resistance profile, b) Plasmid replicon profile and c) virulence gene profile for all colonising *Klebsiella* displayed alongside ST and patient number. Navy indicates resistance/plasmid replicon gene presence and grey resistance/plasmid replicon gene absence. Phylogeny is midpoint rooted with increasing nodes.

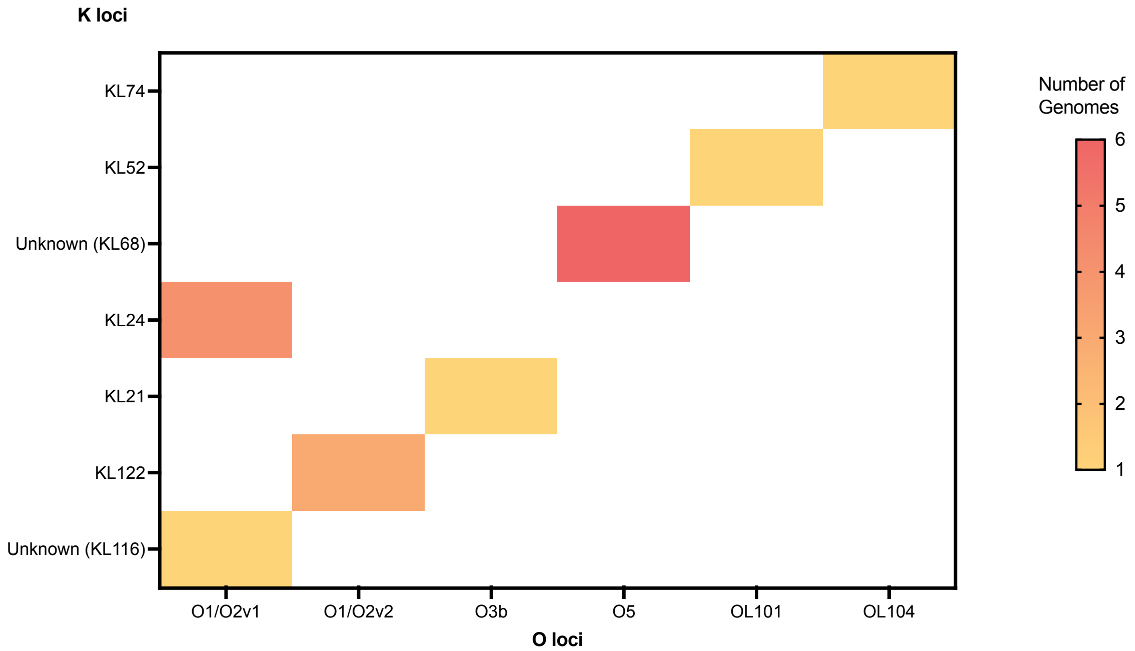


**Fig. S13** K and O loci of all *Klebsiella* isolates as allocated by Kleborate. Number of genomes with the specific K and O loci combination demonstrated by colour gradient.

a)

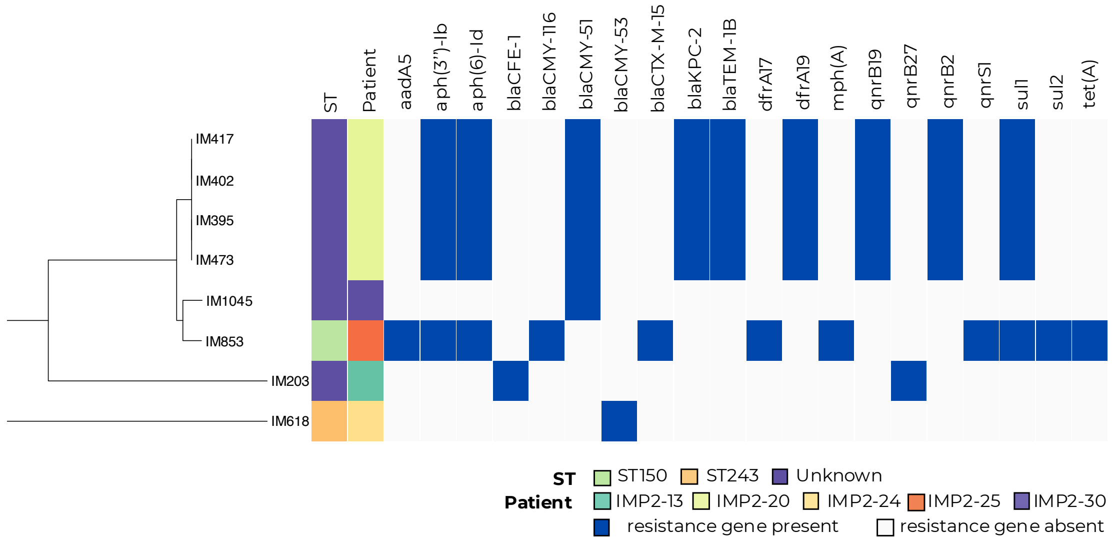


b)

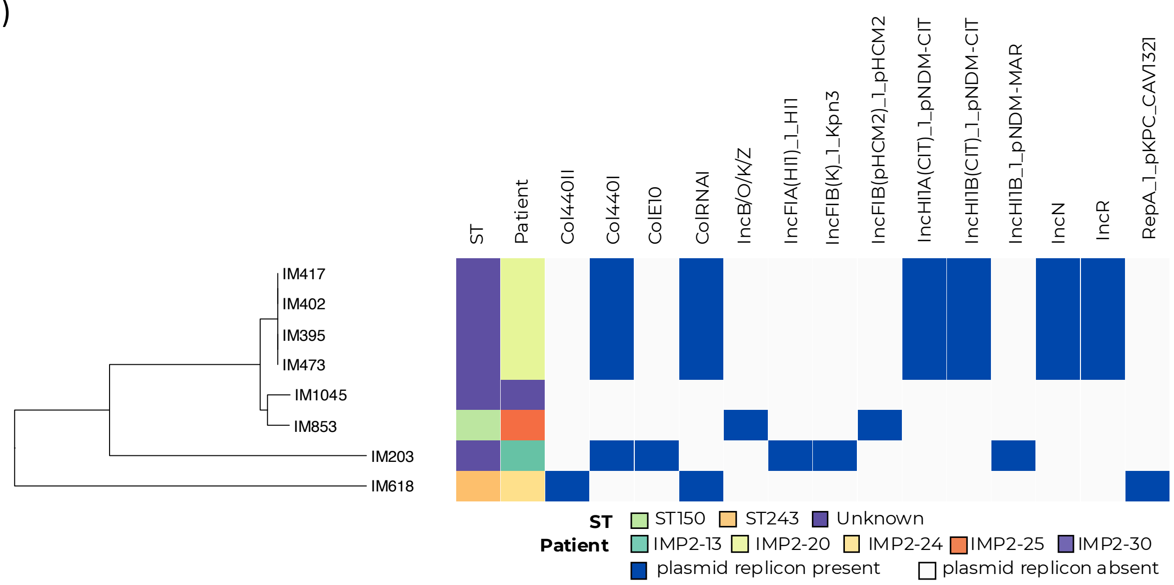


**Fig. S14 a)** Acquired resistance gene profile and b) Plasmid replicon profile for all colonising *C. freundii* displayed next to ST and patient number. There were 8 colonising *Citrobacter freundii* identified in this study, six of which could not be assigned a sequence type. Navy indicates resistance gene/plasmid replicon presence and grey resistance gene/plasmid replicon absence. Phylogeny is midpoint rooted with increasing nodes.

**Table S6** – *Enterobacter cloacae characteristics*

| Isolate | Patient | Day | ST | Patient | Resistance Genes | Plasmid Replicons |
|---|---|---|---|---|---|---|
| IM157 | IMP2-10 | 7 | 45 | IMP2-10 | $bla_{ACT-15}$, *mdf*(A), *oqx*A, *oqx*B | Col440I, ColRNAI, IncR |
| IM212 | IMP2-10 | 8 | 45 | IMP2-10 | $bla_{ACT-15}$, *mdf*(A), *oqx*A, *oqx*B | Col440I, ColRNAI, IncR |
| IM217 | IMP2-10 | 10 | 45 | IMP2-10 | $bla_{ACT-15}$, *mdf*(A), *oqx*A, *oqx*B | Col440I, ColRNAI, IncR |
| IM223 | IMP2-10 | 9 | 45 | IMP2-10 | $bla_{ACT-15}$, *mdf*(A), *oqx*A, *oqx*B | Col440I, ColRNAI, IncR |
| IM1026 | IMP2-37 | 11 | 50 | IMP2-37 | $bla_{ACT-15}$, *fos*A, *oqx*A, *oqx*B | IncFIB(pB171)_1_pB171 |
| IM1031 | IMP2-37 | 14 | 50 | IMP2-37 | $bla_{ACT-15}$, *fos*A, *oqx*A, *oqx*B | - |
| IM1071 | IMP2-37 | 15 | 50 | IMP2-37 | $bla_{ACT-15}$, *fos*A, *oqx*A, *oqx*B | - |
| IM1072 | IMP2-37 | 16 | 50 | IMP2-37 | $bla_{ACT-15}$, *fos*A, *oqx*A, *oqx*B | - |
| IM993 | IMP2-33 | 7 | 50 | IMP2-33 | $bla_{ACT-15}$, *fos*A, *oqx*A, *oqx*B | IncFIB(pENTAS01)_1_pENTAS01, IncFII(pECLA)_1_pECLA |
| IM998 | IMP2-33 | 7 | 50 | IMP2-33 | $bla_{ACT-15}$, *fos*A, *oqx*A, *oqx*B | Col440I, ColRNAI, IncR |
| IM999 | IMP2-33 | 7 | 50 | IMP2-33 | $bla_{ACT-15}$, *fos*A, *oqx*A, *oqx*B | Col440I, ColRNAI, IncR |
| IM585 | IMP2-24 | 9 | 133 | IMP2-24 | $bla_{ACT-7}$, *fos*A, *oqx*A, *oqx*B | - |
| IM670 | IMP2-24 | 16 | 133 | IMP2-24 | $bla_{ACT-7}$, *fos*A, *oqx*A, *oqx*B | IncFIA(HI1)_1_HI1, IncFIB(pECLA)_1_pECLA, IncFII(pECLA)_1_pECLA |
| IM676 | IMP2-24 | 13 | 133 | IMP2-24 | $bla_{ACT-7}$, *fos*A, *oqx*A, *oqx*B | - |
| IM508 | IMP2-21 | 4 | 555 | IMP2-21 | $bla_{ACT-9}$, *fos*A, *oqx*A, *oqx*B | Col440II, Col440I |
| IM227 | IMP2-13 | 8 | 1704 | IMP2-13 | $bla_{ACT-6}$, *mdf*(A), *oqx*A, *oqx*B | Col440II, Col440I |
| IM1065 | IMP2-30 | 27 | - | IMP2-30 | $bla_{ACT-6}$, *fos*A | Col440I, ColRNAI, IncR |
| IM233 | IMP2-13 | 6 | - | IMP2-13 | $bla_{ACT-14}$, *fos*A, *oqx*A, *oqx*B | Col440II, Col440I |

*E. cloacae* did not carry many resistance genes, with all carrying the AmpC ß-lactamase $bla_{ACT}$ and other resistance genes that are normally found on the chromosome of *Enterobacter*.
